# *In vivo* systematic detection of the outcomes of CRISPR/Cas9 mediated DNA repair in skeletal muscle stem cells

**DOI:** 10.1101/2025.08.10.669524

**Authors:** Liangqiang He, Yang Fu, Ziliu Wang, Qin Zhou, Hao Sun, Huating Wang

**Affiliations:** InnoHK Center for Neuromusculoskeletal Restorative Medicine, Hong Kong Science Park, Hong Kong SAR, China; Department of Orthopaedics and Traumatology, Li Ka Shing Institute of Health Sciences, The Chinese University of Hong Kong, Hong Kong, China; Warshel Institute for Computational Biology, Faculty of Medicine, The Chinese University of Hong Kong, Shenzhen, Guangdong, China

## Abstract

CRISPR/Cas9 has revolutionized genome editing with broad therapeutic applications, yet its repair patterns *in vivo* remain poorly understood. Here, we systematically profiled CRISPR/Cas9 editing outcomes at 95 loci using our established CRISPR/Cas9/AAV9-sgRNA system in skeletal muscle stem cells (MuSCs). Through comprehensive characterization of the repair outcomes, our findings demonstrate that the general rules governing CRISPR/Cas9-mediated editing *in vivo* largely align with those observed *in vitro* but with reduced editing precision. Additional to the anticipated small editing indels such as MMEJ mediated deletions and NHEJ mediated templated insertions, we uncovered a prevalent occurrence of large on-target modifications, including large deletions (LDs) characterized by microhomology (MH) and large insertions (LIs). Notably, the LIs comprise not only exogenous AAV vector integrations but also endogenous genomic DNA fragments (Endo-LIs). Endo-LIs preferentially originate from active genomic regions, with their integration shaped by three-dimensional chromatin architecture. By disrupting key components of the NHEJ and MMEJ repair pathways *in vivo*, we identified their distinct roles in regulating the large on-target modifications. Together, our work for the first time systematically profiles the CRISPR/Cas9 repair outcomes *in vivo* and offers valuable guidance for improving the safety of CRISPR/Cas9-based gene therapies.

## Introduction

CRISPR/Cas9 (Clustered regularly interspaced short palindromic repeats and CRISPR-associated protein 9) represents a groundbreaking genome editing technology that has revolutionized molecular biology and the development of potential therapies for human genetic disorders^1,2^. It initiates targeted DNA modifications by the recognition of target sequence through a single guide RNA (sgRNA). The sgRNA first binds to the Cas9 protein, forming a ribonucleoprotein complex to locate its target next to a protospacer adjacent motif (PAM), typically “NGG” in the case of *Streptococcus pyogenes* Cas9 (SpCas9). Upon PAM recognition, the sgRNA hybridizes with the complementary DNA strand, forming an R-loop structure, where the non-target strand remains single-stranded. The Cas9 protein harbors two nuclease domains, HNH and RuvC, which cleave the target and non-target DNA strands, respectively^1,2^. Typically, both cleavages occur 3 bp upstream of the PAM motif, leading to a blunt ended double-strand break (DSB). However, accumulating evidence suggests that while the HNH domain cleaves precisely, the RuvC domain exhibits flexibility, cutting at positions –3 bp, –4 bp, –5 bp, or even further upstream, leading to the formation of both blunt and 5’ staggered ends^3,4^.

In eukaryotic cells, CRISPR/Cas9 generated DSBs are primarily repaired through three competing pathways: homology-directed repair (HDR), non-homologous end joining (NHEJ) and microhomology-mediated end joining (MMEJ)^5^. HDR is a high-fidelity repair mechanism that uses a homologous DNA template to precisely repair the break. However, HDR is primarily active during the S and G2 phases of the cell cycle but significantly inactive in non-dividing or terminally differentiated cells. In contrast, NHEJ is the predominant repair pathway used in most cells and is characterized by its rapid kinetics. This pathway acts without the need for a homologous template and typically introduces small insertions or deletions (indels) at the break site, normally within 2 bp^6^. Although NHEJ is generally considered error-prone, emerging evidence from cultured cells *in vitro* or massively synthetic target sequences suggests that it can mediate precise and predictable repair of Cas9-induced DSBs^3,4,7,8^. Once a DSB forms, NHEJ directly ligates the two broken DNA ends without modifications, leading to repeated cycles of Cas9 cleavage until an indel occurs to prevent further sgRNA target recognition^5,9^. When staggered ends with 5’ overhangs arise due to the flexible cutting of the RuvC domain, the NHEJ pathway fills in the gaps and joins the two ends together, resulting in insertions that are templated by the sequence upstream of the cut site. In single sgRNA-induced editing, the most consistently predictable outcome is the insertion of a single nucleotide identical to the nucleotide located -4 bp upstream of the PAM motif^3,8^. Accordingly, in dual sgRNAs-induced fragment excision, the two DSBs are preferentially ligated accurately by NHEJ, resulting in precise deletions or templated insertions^7^. Beyond HDR and NHEJ, alternative repair pathways, such as MMEJ, or Polymerase Theta-mediated end-Joining (TMEJ) as recently described^10,11^, can also contribute to DSB repair. Like NHEJ, MMEJ does not require a template for repairing the damaged DNA but utilizes short microhomologies (MH) present near the broken DNA ends to operate the DSBs^5,6^. MMEJ is normally responsible for generating small deletions (≥ 3 bp) within 10 bp^6,12^, but growing studies suggest that MMEJ can also lead to the occurrence of large deletions (> 50 bp)^10,13,14^.

The general indel patterns induced by CRISPR/Cas9 have been extensively studied *in vitro* especially using a synthetic reporter-based system containing a sgRNA expression cassette together with its PAM-containing target sequence flanked by common PCR priming sites on both sides^8,15–17^. High-throughput profiling of repair outcomes leveraging this system indicates that the indels resulted from end-joining repair of these DSBs are predictable and significantly influenced by the sequence context surrounding the targeted site^8,15–17^. Based on these findings, several sophisticated machine learning models have been developed to predict the predominant indels of Cas9-induced DSBs. Besides, the local chromatin structure also impacts Cas9 generated indel patterns^18^. Notably, NHEJ-mediated repair of Cas9-induced DSBs is more efficient in euchromatin, whereas MMEJ is preferentially active in specific heterochromatic regions. Mechanically, NHEJ is typically initiated by the Ku70–Ku80 heterodimer following DSB formation, which binds to broken DNA ends to prevent resection^5,6^. This complex subsequently recruits DNA-PKcs and DNA Ligase IV (Lig4) to facilitate end ligation^5,6^. In contrast, HDR and MMEJ require end resection by the Mre11–Rad50–Nbn (MRN) complex and its cofactor CtIP to expose MH. Strategies to regulate the DSB repair pathway choice by modulating the activities of these key components have been developed to facilitate more efficient and precise editing. In CRISPR/Cas9-related *ex vivo* gene therapy, a common strategy is to enhance HDR levels by simultaneously suppressing MMEJ and NHEJ activity using specific inhibitors targeting key factors of these pathways^5,6^.

Other than the predictable small indels, recent studies have identified undesirable editing byproducts: complex on-target large gene modifications, which raise critical concerns about the safety in clinical applications of CRISPR/Cas9^13,19,20^. Gross genomic modifications can occur at the target site, including large deletions (LDs), large insertions (LIs)—especially the integration of exogenous vector DNA—and even chromosomal structural rearrangements and chromosomal loss^21–23^. These unintended alterations arise from the cellular repair triggered by Cas9-induced DSBs thus cannot be avoided by using high-fidelity Cas9 variants. Typically, these byproducts occur at low frequencies, making it unclear whether they are merely incidental to the editing process or represent a more widespread phenomenon. Overall, the current understanding of on-target large genomic modifications remains limited. It is shown that MHs are enriched at LD breakpoint junctions, and LD frequency increases in NHEJ-deficient cells but declines in cells with impaired MMEJ activity, implicating MMEJ in LD formation^13,14^. On the other hand, LIs frequently arise from the integration of exogenous vector fragments into DSBs within the host genome. The mechanisms underlying vector integration remain unclear; however, abolishing either NHEJ or MMEJ pathways can inhibit random integration, indicating that both pathways may contribute to CRISPR/Cas9-induced large exogenous vector insertions^24,25^. In addition, recent studies have revealed that LIs at CRISPR/Cas9-induced DSBs can also originate from endogenous genomic sequences^23,26^. Comprehensive characterization of these endogenous LIs is currently lacking, and their mechanisms of formation remain poorly understood.

CRISPR-based gene editing has made remarkable progress in treating diverse human genetic disorders^1,27^. In particular, *in vivo* genome editing, which delivers CRISPR machinery directly to affected tissues *via* localized or systemic administration, offers the promise of targeting a broader spectrum of diseases. Advanced delivery platforms leverage both non-viral and viral based vectors, with adeno-associated virus (AAV) emerging as the leading choice for *in vivo* gene therapy applications due to its tissue-specific tropism, sustained transgene expression, and favorable safety profile^28^. While *in vitro* CRISPR/Cas9-induced DNA repair has been extensively studied, a systematic characterization of its performance *in vivo* remains lacking. Given that the DNA repair process may be modulated by cellular context and tissue-specific factors^5^—which are still poorly understood—it remains uncertain whether the repair patterns observed *in vitro* fully recapitulate those *in vivo*. Moreover, the presence of large on-target genomic modifications needs further investigation. Although the incidence of such large chromosomal aberrations could be rare *in vivo*, even a small number of edited cells harboring deleterious mutations could clonally expand and lead to serious abnormalities. For instance, while AAV predominantly remains episomal, AAV vector integrated into DSBs within the host genome can be detected after *in vivo* administration^9,29,30^. Therefore, systematically analyzing these unwanted byproducts and elucidating their formation mechanisms to minimize their occurrence *in vivo* is crucial for improving the safety of CRISPR/Cas9 based therapies.

In our previous work, we established a CRISPR/Cas9/AAV9-sgRNA platform for *in vivo* editing in skeletal muscle stem cells (MuSCs) characterized by paired-box gene 7 (PAX7) expression^31,32^. This platform presented high efficiency in introducing mutagenesis at multiple genomic loci in MuSCs, enabling it as a robust tool for studying CRISPR/Cas9 mediated editing outcomes *in vivo*^31,33–35^. Here, leveraging this system, we systematically edited 95 loci in MuSCs *in vivo* using single or dual sgRNAs. Through targeted deep sequencing, we performed in-depth characterization of CRISPR/Cas9-induced repair profiles to decipher the indel patterns and assess large on-target modifications. Our results indicate that CRISPR/Cas9-induced indel patterns *in vivo* largely align with those observed *in vitro*, although exhibiting reduced precision. Moreover, we uncovered prevalent large on-target modifications *in vivo*, including LDs and LIs. Notably, the LIs comprise not only exogenous AAV vector integrations but also endogenous genomic DNA fragments. These endogenous fragments are preferentially derived from active genomic regions, and the insertion is influenced by the 3D chromatin architecture. Finally, through knocking out key factors involved in NHEJ and MMEJ *in vivo*, we revealed their differential regulatory effects on these large on-target modifications. Collectively, our work for the first time systematically uncovers the CRISPR/Cas9 repair outcomes *in vivo* and offers valuable guidance for enhancing the safety of CRISPR/Cas9-based *in vivo* gene therapies.

## Results

### *In vivo* mapping of CRISPR/Cas9 induced indel profiles in MuSCs using single-sgRNAs

To uncover the patterns of indel profiles induced by CRISPR/Cas9 *in vivo*, we took advantage of our previously developed CRISPR/Cas9/AAV9-gRNA editing system to conduct a multi-loci profiling of the repair outcomes in MuSCs *in vivo*^31,32^ (Fig. 1a). In brief, a muscle-specific Cas9 knockin mouse line was generated by crossing Pax7^Cre^ with Cre-dependent Cas9 knockin mice (referred to as Pax7^Cas9^ mouse)^31–35^. Skeletal muscle-tropic AAV9 virus containing locus specific single or dual sgRNA was intramuscularly (IM) injected into the Pax7^Cas9^ mouse at postnatal day 10 (P10), followed by MuSC isolation four weeks after injection. This strategy has been successfully applied in our prior studies to introduce mutagenesis at multiple target loci in MuSCs *in vivo*^31–35^. 44 single sgRNAs were designed to target distinct genomic sites, designated numerically from S1 to S44 (Supplementary Table 1). These sites covered coding sequences (CDS) and promoter regions of seven transcription factors, including *Myc*, *Bcl6*, *Myod1*, *Rora*, *Runx1*, *Pknox2* and *Ctcf*. Additionally, 13 were designed to target potential enhancer regions, with three located near *Pax7* gene and another three near *Runx1*. To further increase the diversity and complexity of the targeted sites, we also designed three sgRNAs targeting non-regulatory intergenic regions (Fig. 1b, Extended Data Fig. 1a and Supplementary Table 1). Two biological replicates were included for each target site. Sequences encompassing each sgRNA targeting site were amplified and subjected to unbiased next generation sequencing (NGS) for precise characterization of editing efficiency and indel profiles (Fig. 1a). The sequenced data was computed by Sequence Interrogation and Quantification (SIQ)^36^, yielding over 10,000 mappable reads per sample (Extended Data Fig. 1b and Supplementary Tables 2 and 3), and generating a comprehensive indel profile at each target site according to mutation characteristics^36^. To ensure a consistent basis for monitoring the distribution of the indel profiles across amplicons of diverse sizes, only indels overlapping within a 60 bp window centered on the canonical cut site (3 bp upstream of the NGG PAM sequence) were analyzed (Fig. 1c). We first examined the editing efficiency by calculating the total indel frequency (TIF) and found it varied across different loci (ranging from 0.7% to 88.2%, with an average of 32.5%) (Fig. 1d). A strong similarity between the two replicates of the same locus was detected (Pearson’s R: 0.951) (Extended Data Fig. 1c). As expected, the TIF generally correlated positively with active chromatin regions featured by ATAC-seq, H3K4me3, H3K27ac and H3K4me1 ChIP-seq signals but not repressive H3K27me3 ChIP-seq signals^6,18^ (Fig. 1e).

**Figure 1.**
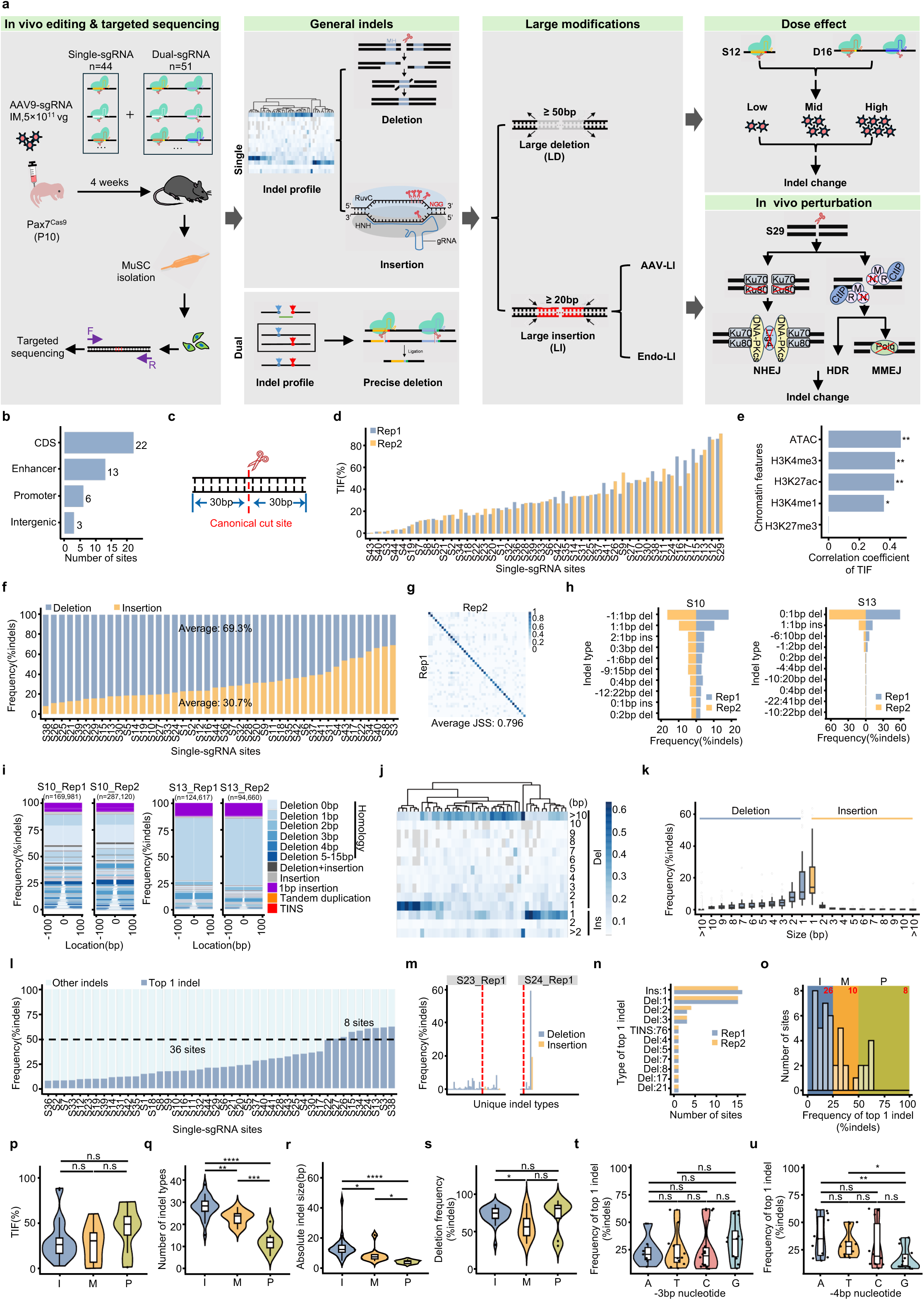
*In vivo* mapping of CRISPR/Cas9 induced indel profiles in MuSCs using single-sgRNAs. (**a**) Overview of the experimental design/analysis for the study. (**b**) The number of target sites with the indicated genomic annotations. (**c**) Schematic illustration of the 60 bp editing window centered on the canonical cut site indicated as a red dashed line. Only indels overlapping with this region were included for analysis. (**d**) TIF for the 44 targets (each with 2 biological replicates). (**e**) Spearman’s correlation between TIF and indicated chromatin features. (**f**) Frequencies of deletions and insertions in total indels, averaged from two biological replicates per target. The average frequencies of deletions and insertions across all targets are shown. (**g**) Heatmap of Jaccard similarity scores (JSS) of Rep 1 and Rep2 for targets S1 to S44. Each square indicates the JSS between the top ten most frequent indels of the corresponding target, while the diagonal highlights scores for identical targets. The average JSS of paired replicates across all targets is shown. (**h**) Frequency of the top ten indel types in each replicate of S10 (Left) and S13 (Right). The y-axis displays the indel identity (start coordinate relative to the breakpoint): (indel size and type). (**i**) Visualized tornado plots showing the frequency of indels of the two S10 (Left) and S13 (Right) replicates. The indels are sorted by deletion size, and the x-axis displays genomic positions relative to the canonical Cas9 cut site (position 0). White space in the center indicates deleted regions. The intensity of blue coloring corresponds to the degree of MH utilized in the formation of deletions. (**j**) Heatmap illustrating the frequencies of indels categorized by size across 44 targets. The targets are clustered using the “complete” hierarchical clustering. (**k**) Boxplot illustrating the distribution of indels by size across targets. (**l**) Frequency of the top 1 and other indels in total indels for each target. The number of sites with top 1 indel frequency > (8) or ≤ (36) 50% is shown. (**m**) Indel profiles of an imprecise (S23, Left) and a precise (S24, Right) target. The canonical cut site is marked by a red dashed line. (**n**) The number of sites with the indicated type of top 1 indels in Rep1(blue) and Rep2 (orange) across all target sites. (**o**) Distribution of the frequency of top 1 indels across all target sites. Imprecise(I), middle(M), precise(P) groups are indicated in blue, orange and green. Inset numbers marked by red indicate the count of target sites in each group. (**p**-**s**) The TIF (**p**), number of indel types (**q**), absolute indel size (**r**) and frequency of deletions (**s**) in I, M and P groups, respectively. (**t**-**u**) Frequency of top1 indel in total indels at sites with A, T, C, G at -3 bp (**t**) or -4 bp (**u**) upstream of the PAM motif. Only indels with frequency > 0.5% of the total indels were considered for (**m**) and (**q**). Statistical significance for (**p**) - (**u**) was calculated using a t-test. *p < 0.05, ** p < 0.01, ***p < 0.001, ****p < 0.0001 and ns, no significance.

We next examined the indel profiles to uncover the general patterns of repair outcomes driven by single-sgRNA-directed Cas9 activity. The indels were quantified based on their mutation types (deletions or insertions). Of note, complex repair outcomes with simultaneous insertions and deletions were classified as insertions^37^. Interestingly, we found that deletions predominated the overall indel landscape, accounting for an average of 69.3% of the total indels (Fig. 1f). We then assessed if the editing is reproducible as observed *in vitro*^3,8,38,39^. A Jaccard similarity score (JSS) was calculated for the top ten most frequent indels (Supplementary Table 4), which represented an average of 59.2% of all indels at each site (Extended Data Fig. 1d-1e), and high similarity (0.796) was found between the two replicates while samples from different sites showed minimal similarity (Fig. 1g and Supplementary Table 4), suggesting the indel formation is not random and target specific. As exemplified at the S10 and S13 (Fig. 1h) loci, the top ten indel types—indicted by the type of the indel, the coordinate of the start site relative to the break point, the indel length, and the frequency— exhibited remarkable similarity; for example, the top 1 most frequent indel type in both replicates of S10 is a 1 bp deletion occurring 1 bp upstream of the breakpoint (-1:1bp del). To further dissect the reproducibility of the editing outcomes, we generated a visualized plot of the overall mutational outcomes using the SIQ software by categorizing the indel profiles based on common genetic variations, such as insertions, deletions, deletions with insertions, tandem duplications, and deletions with templated insertions (TINS). TINS, in particular, has been identified as an evolutionarily conserved mutational signature associated with DNA polymerase theta (Pol θ)-mediated end joining (TMEJ)^36^. Our results revealed that the overall indel spectrum was also similar between the two independent biological replicates (Fig. 1i), and this is consistent with findings from *in vitro* studies^3,8,38,39^.

Further unsupervised hierarchical clustering based on the frequencies of the identified indel classes revealed four major groups of sites displaying distinct indel signatures, including sites that preferentially exhibited 1 bp deletions, deletions > 10 bp, 1 bp insertions, and > 10bp deletions together with > 2 bp insertions (Fig. 1j). Overall, 1 bp insertions and deletions were the predominant mutational types, larger insertions and deletions occurred infrequently and the frequencies declined with increasing size (Fig. 1k); nevertheless, insertions and deletions larger than 10 bp were also noticed.

It has been reported that the level of editing precision can be assessed by the recurrence of specific indels at each target^38^, we thus ranked all sites based on the prevalence of the top 1 most frequent indel. Our result showed that only eight sites (8/44) exhibited a specific mutation type with a probability > 50% (Fig. 1l). Some sites displayed a remarkably diverse range of indel types with moderate frequency. For example, at the S23 site (Fig. 1m, Left), we detected up to 37 different types without a prominent one. In contrast, others such as the S24 site (Fig. 1m, Right) showed a strong bias toward a single predominant indel, accompanied by only a few additional types, suggesting that the editing precision varied from site to site. Consistent with the highest frequency observed (Fig. 1k), 1 bp insertions and 1 bp deletions presented as the top 1 indel for 31 sites (Fig. 1n). Nevertheless, targets favoring longer deletions up to 17 or 21 bp were also observed.

To delve into the factors that may be associated with editing precision, 44 target loci were divided into three groups^38^: imprecise (I, the frequency of top 1 indel ≤ 0.25), medium precision (M, 0.25 < the frequency of top 1 indel ≤ 0.5) and precise (P, 0.5 < the frequency of top 1 indel ≤ 1) (Fig. 1o). Based on this classification, over half of the sites (26/44) were categorized into the imprecise group, which is significantly higher than previously observed *in vitro*^38^, suggesting Cas9-induced indel profiles *in vivo* may be more complex. Also, the TIF showed no significant difference among the three groups (Fig. 1p). As expected, editing precision exhibited a negative correlation with the number of indel types, with the average number of indel types at imprecise sites (28.1) two times greater than that at precise sites (12.6) (Fig. 1q). A similar pattern was observed with the absolute indel size, where precise sites were more prone to producing short indels (Fig. 1r). Moreover, the group I exhibited a significantly higher portion of deletions compared to group M but not group P (Fig. 1s). Previous studies have demonstrated that the nucleotide composition flanking the cut site—particularly at the position immediately downstream (-3 bp relative to PAM) and upstream (-4 bp)—can significantly influence DNA repair outcome^3,40^, we thus examined whether nucleotide identity at these specific positions correlates with editing precision. At the -3 bp position, the presence of G was associated with an elevated but not significant mean frequency of the top 1 indels compared to other nucleotides (Fig. 1t). Conversely, the -4 bp G was associated with a lower mean frequency of predominant indels especially compared to A and T (Fig. 1u). These nucleotide-dependent effects indicate that local DNA context surrounding the cleavage site may modulate editing precision *in vivo*.

Finally, we investigated the impact of chromatin states on editing precision, as recent studies have underscored the significance of chromatin context in determining the choice of DSB repair pathways *in vitro*^6,18^. The 44 sites were categorized into two groups based on their location in either peak or non-peak regions of various chromatin features including H3K4me3, H3K4me1, H3K27ac, H3K27me3 and chromatin openness determined by ATAC-seq (Extended Data Fig. 2a). By comparing the types of the top 1 indel, we found no obvious differences between the two categories, as 1 bp indels consistently predominated across all conditions (Extended Data Fig. 2b). This suggests that the chromatin environment has no apparent effect on *in vivo* editing precision. Altogether, these observations suggest that CRISPR/Cas9 induced indels in MuSCs *in vivo* displays high producibility and low editing precision that is affected by local DNA context.

### Extensive use of microhomology in CRISPR/Cas9 induced deletions in MuSCs *in vivo*

Considering that deletions constituted the majority of indels (Fig. 1f), we subsequently investigated the underlying patterns of deletions. It is known that MH-mediated alignment of broken ends is commonly employed in CRISPR/Cas9-induced deletions^8,38^(Fig. 2a), we therefore systematically evaluated the presence of MH at the breakpoint junctions. Our analysis indicated that 73.1% of the deletions were indeed characterized by the presence of MH (Fig. 2b). The length of the MH varied from 1 to 11 bp, with the frequency of associated deletions gradually decreasing as MH size increased (Fig. 2c and Extended Data Fig. 3a). Overall, short MH tracks (1-4 bp) constituted most MH-mediated deletions (approximately 95%) (Extended Data Fig. 3b); Deletions utilizing 1 bp MH occurred most frequently, accounting for 35.68% of all deletions and were detectable at all sites. In contrast, longer MHs (MH > 4 bp) were observed at a significantly lower frequency, with 11 bp MH deletion detected at only two sites (S21 and S23) (Extended Data Fig. 3a-3c). The frequency of MH deletions shorter than 7 bp was substantially higher than the background expectation (Fig. 2c). Notably, we found that longer MH sizes were correlated with larger average deletion sizes (Fig. 2d and Extended Data Fig. 3c). Moreover, MH-mediated deletions exhibited distinct characteristics compared to non-MH deletions. First, the average size of MH deletions was larger (15 vs. 6 bp) (Fig. 2e-2f). Further unsupervised hierarchical clustering revealed that the length of MH deletions peaked at both 1 bp and 10-20 bp, with 41% exceeding 10 bp (Fig. 2g). Conversely, non-MH-mediated deletions produced smaller deletions, with approximately half (48%) being 1-2 bp deletions (Fig. 2h).

**Figure 2.**
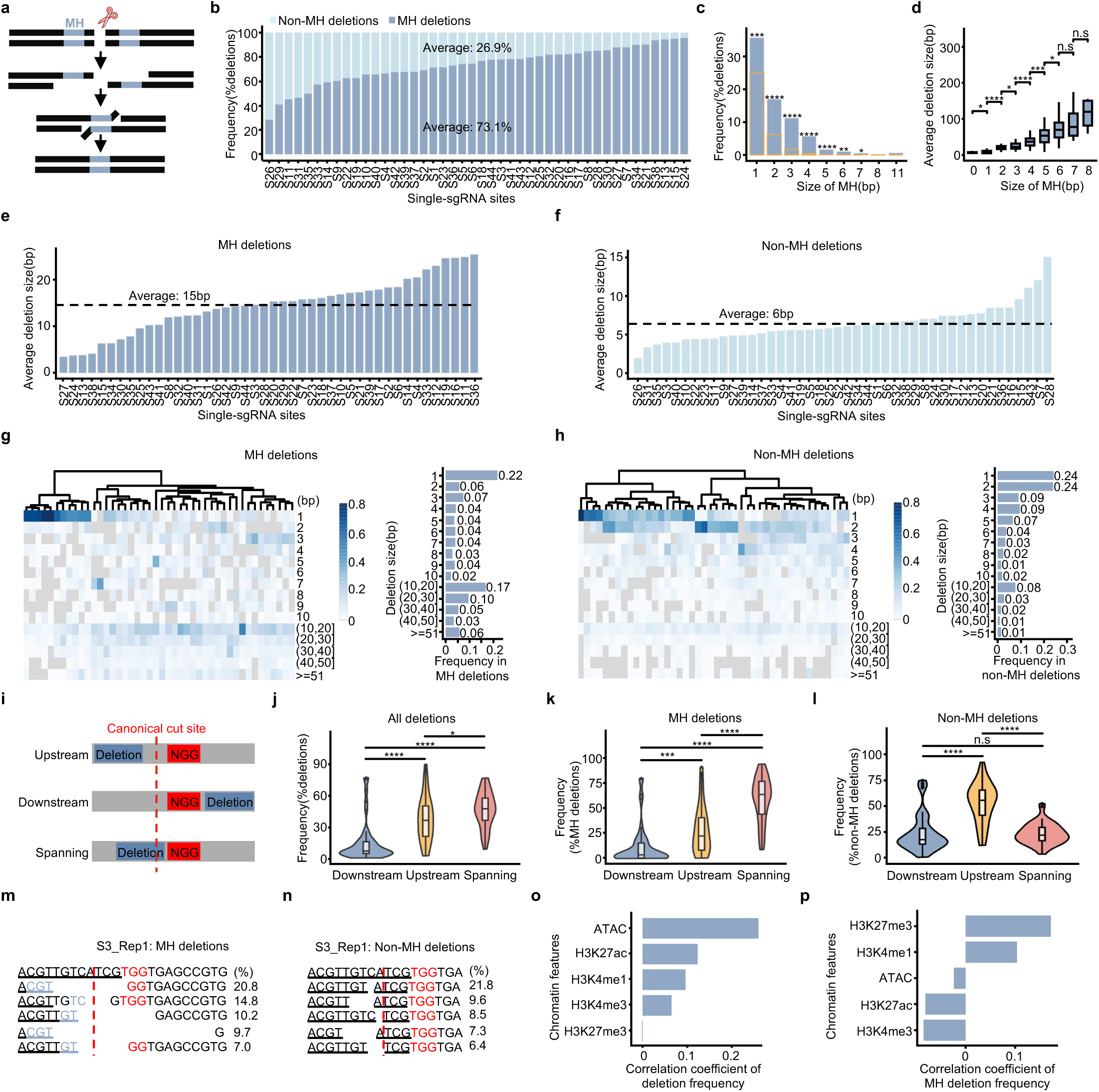
Extensive use of microhomology in CRISPR/Cas9 induced deletions in MuSCs *in vivo*. (**a**) Schematic illustrating MH dependent deletions in CRISPR/Cas9 induced DSB repair. MH tracts (blue) flanking DSB facilitate the alignment of the broken ends. The unpaired overhang is cleaved by endonucleases, followed by DNA polymerases mediated gap filling and DNA ligases mediated breakpoint ligation. (**b**) Frequency of MH and non-MH mediated deletions in all deletions for each site. The average frequencies of MH and non-MH mediated deletions across all sites are shown. (**c**) Frequency of deletions with MH of varying sizes in all deletions. The orange bar represents the expected frequency for each MH size. (**d**) Average deletion size for deletions categorized by MH sizes. (**e**-**f**) Average deletion size for MH (**e**) and non-MH (**f**) mediated deletions for each site. MH and non-MH mediated average deletion sizes across all sites are shown by the dash line. (**g**) Left: Heatmaps illustrating the frequency of MH mediated deletions (a proportion of total MH mediated deletions) categorized by size across 44 targets. The targets are clustered using the “complete” hierarchical clustering. Right: The average frequency (indicated by inset numbers) across all targets for each category with varying deletion size in MH mediated deletions. (**h**) The above analysis in non-MH mediated deletions. (**i**) Schematic diagram illustrating deletion directions. The NGG PAM motif is highlighted by red background. (**j-l**) Violin plots illustrating the frequency of all deletions (**j**), MH (**k**), and non-MH (**l**) mediated deletions categorized by direction. (**m-n**) An example of target site S3 illustrating top five most frequent MH (**m**) and non-MH (**n**) mediated deletions. The frequency is presented as percentage in all MH or non-MH deletions. The canonical cut site is indicated by a red dashed line and the NGG PAM motif is highlighted in red. The guide sequence is underlined. (**o**-**p**) Spearman’s correlation between deletion frequency (**o**) or MH frequency in deletions (**p**) and indicated chromatin features. Statistical significance was calculated using a t-test for (**d**), (**j**) - (**l**) and a one-tailed t-test (alternative = ’greater’) for (**c**). *p < 0.05, **p < 0.01, ***p < 0.001, ****p ≤ 0.0001 and ns, no significance.

It is reported that CRISPR/Cas9-induced deletions are asymmetric and occur unidirectionally downstream of the breakpoint^8^, we thus analyzed the positioning patterns of the deletions relative to the canonical cut site by classifying deletions into three groups: downstream deletions occurring within the PAM-containing flank, upstream deletions occurring in the opposite flank, and spanning deletions across the canonical cut site (Fig. 2i). Our results showed that among all the deletion events, on average, 46.7% spanned the cleavage site, followed by 37.9% upstream and 15.4% downstream deletions (Fig. 2j). MH-mediated deletions exhibited a similar pattern (59.0%, 28.9% and 12.1% spanning, upstream and downstream deletions) (Fig. 2k and Extended Data Fig. 3d), indicating that most CRISPR/Cas9-induced *in vivo* deletions were not asymmetric, which contrasts with the *in vitro* findings^8^. Non-MH deletions also exhibited unidirectional distribution but interestingly had a preference for upstream rather than downstream deletions (Fig. 2l), which is also different from the *in vitro* finding^8^. For instance, at the S3 locus, the top five MH deletions (accounting for 62.5% of total MH deletions) all spanned the cut sites (Fig. 2m). In contrast, the top five non-MH deletions (representing 53.6% of total non-MH deletions) primarily occurred upstream of the cut sites (Fig. 2n).

Finally, we examined the influence of chromatin context on deletion events. While deletions tended to occur in open euchromatin regions characterized by ATAC and H3K27ac signals (Fig. 2o), MH-deletions exhibited a potential association with heterochromatin contexts marked by H3K27me3 modification (Fig. 2p); although lacking statistical significance, possibly due to our limited sample size, this observation is consistent with previous *in vitro* findings^6,18^. Altogether the above results indicate that MH is prevalent in CRISPR/Cas9-induced deletions in MuSCs *in vivo* and MH mediated deletions display distinct features from non-MH mediated deletions.

### CRISPR/Cas9-mediated short insertions in MuSCs *in vivo* are templated and predictable

Next, we took a close examination of the insertions detected in *vivo*. *In vitro* studies have demonstrated that Cas9-mediated short insertions are precise and predictable even without donor templates, reflecting the error-free nature of NHEJ in the repair of Cas9 generated DSBs^3,4,8,41^. Specifically, the 1 bp insertion is predominantly templated by the nucleotide located upstream of the cleavage site (–4 bp upstream of the NGG PAM motif) (so called templated insertions, TIs, excluding the TINS where deletions and templated insertions appear simultaneously), thus the inserted sequence tends to be identical to the -4 bp nucleotide (Fig. 3a). We found the frequency of TIs in MuSCs *in vivo* varied significantly across the 44 sites, ranging from 15.4% to 100%, with an average of 74.1% (Fig. 3b). Moreover, the frequency declined sharply as the insertion size increased with 1 bp TI being the predominant type (96.6%) and those larger than 3 bp rarely detected (0.1%) (Fig. 3c-3d). Furthermore, the frequency of TIs with short size (1, 2 or 3 bp) was significantly higher than the background expectation (Fig. 3e).

**Figure 3.**
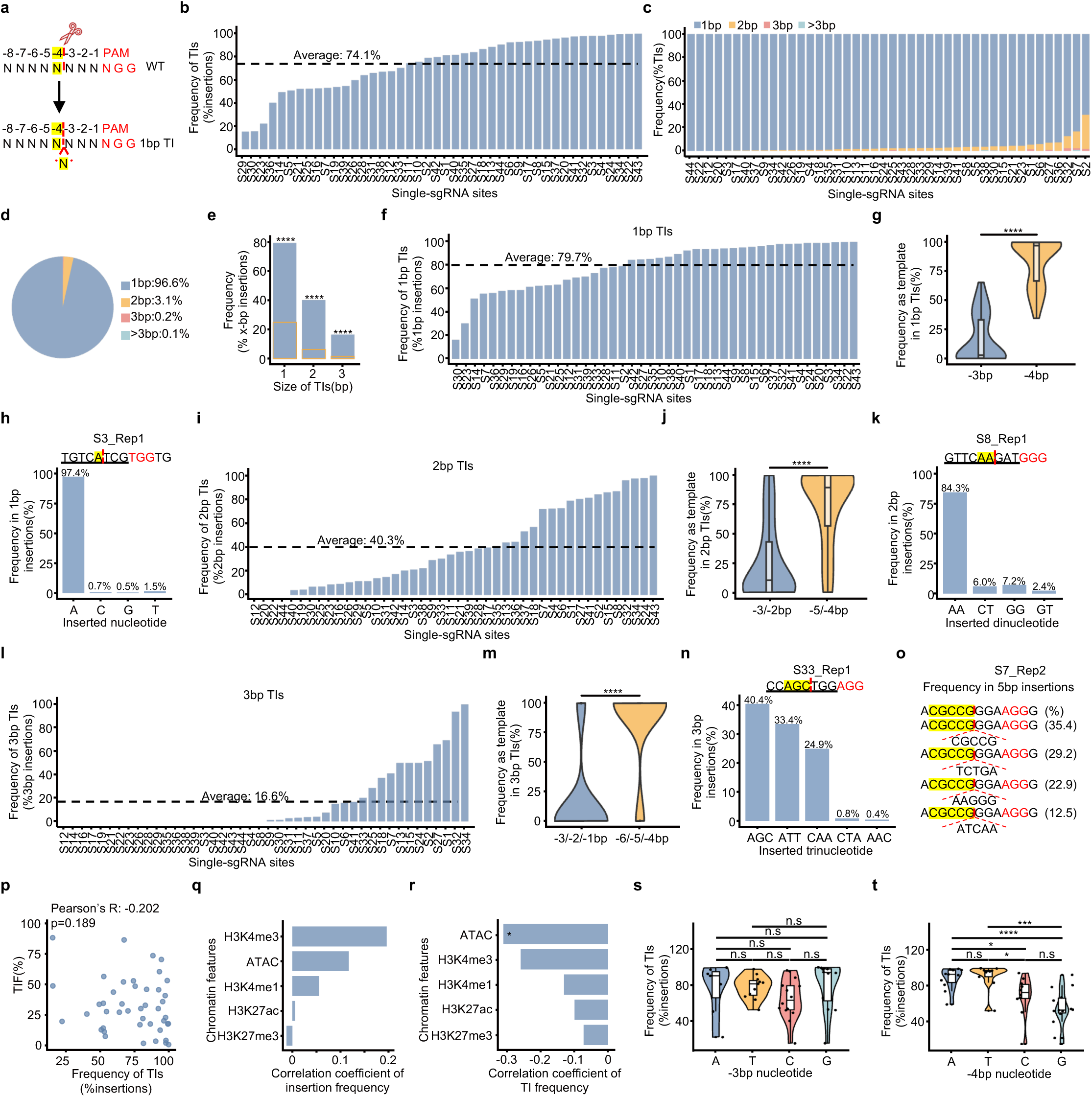
CRISPR/Cas9-mediated short insertions in MuSCs *in vivo* are templated and predictable. (**a**) Schematic illustration of 1 bp TI. The -4 bp position nucleotide functioning as the template is marked as yellow. (**b**) Frequency of TIs in all insertions at the canonical cut site across 44 single-sgRNA target sites. The average frequency of TIs across all sites is shown by the dash line. (**c**) Frequency of TIs with varying sizes in all TIs for each target site. (**d**) Pie chart showing the average frequency of TIs of varying sizes. (**e**) Frequency of 1 bp, 2 bp and 3 bp TIs, with the orange bar representing the expected frequency for each TI category. (**f**) Frequency of 1 bp TIs in all 1 bp insertions for each target site. The average frequency of 1 bp TIs across all sites is shown by the dash line. (**g**) Relative frequency of the upstream -3 bp or -4 bp nucleotide as template in 1 bp TIs. (**h**) An example of 1 bp TIs on target S3. The bar plot illustrates the frequencies of inserted nucleotide being A, T, C, G in 1 bp insertions. The nucleotide at -4 bp upstream of the PAM motif is highlighted in yellow. The canonical cut site is indicated by a red dashed line and the PAM motif is marked in red. (**i**) Frequency of 2 bp TIs in all 2 bp insertions for each target site. The average frequency of 2 bp TIs across all sites is shown by the dash line. (**j**) Relative frequency of the -3 to -2 bp or -5 to -4 bp dinucleotides as templates in 2 bp TIs. (**k**) An example of 2 bp TIs on S8. The bar plot illustrates the frequencies of all detected dinucleotides in 2 bp insertions. The dinucleotides at -5 to -4 bp upstream of the PAM motif are highlighted in yellow. (**l**) Frequency of 3 bp TIs in all 3 bp insertions for each target site. The average frequency of 3 bp TIs across all sites is shown by the dash line. (**m**) Relative frequency of the -3 to -1 bp or -6 to -4 bp trinucleotides as templates in 3 bp TIs. (**n**) An example of 3 bp TIs on S33. The trinucleotides at -6 to -4 bp upstream of the PAM motif are highlighted in yellow. The bar plot illustrates the frequencies of all detected trinucleotides in 3 bp insertions at the canonical cut site for S33. (**o**) An example of > 3 bp TIs on S7. The nucleotides at -8 bp to -4 bp upstream of the PAM motif are highlighted in yellow. The frequency is presented as percentage in 5 bp insertions. (**p**) Correlation between TIF and TI frequency in all insertions. (**q**-**r**) Spearman’s correlation between insertion frequency (**q**) or TI frequency (**r**) and indicated chromatin features. (**s**-**t**) Frequency of TIs in all insertions at target sites with A, T, C, G at -3 bp (**s**) or -4 bp (**t**) upstream of the PAM motif. Statistical significance was calculated using a t-test for (**g**), (**j**), (**m**), (**s**) and (**t**) and a one-tailed t-test (alternative = ’greater’) for (**e**). *p < 0.05, *** p < 0.001, ****p < 0.0001 and ns, no significance.

As the predominant type, 1 bp TIs were detected at all 44 sites, accounting for 79.7% of the total 1 bp insertions (Fig. 3f). As expected, -4 bp nucleotides was more frequently (83.2% vs. 16.8%) used as the template compared to -3 bp nucleotides (Fig. 3g), indicating asymmetric templating as previously reported^3,4,8,41^. As a result, the 1 bp insertions tended to be identical to the -4 bp nucleotides; for example, when the base at the -4 bp is A at S3 site, the inserted nucleotide is most likely to be A; 1 bp TIs using the -4 bp A as the template accounted for over 97.4% of all 1 bp insertions (Fig. 3h). As for 2 bp TIs, although they represented only 40.3% of all 2 bp insertions (Fig. 3i)—substantially lower than that of 1 bp TIs (Fig. 3f)—a similar asymmetric templating was observed; the dinucleotides located upstream (-5/-4 bp relative to PAM) of the cut site were more frequently (74.5% vs. 25.5%) used as the repair template compared to the downstream (-3/-2 bp relative to PAM) sequences (Fig. 3j). For example, as shown at the S8 site, 84.3% of all 2 bp insertions used the -5 to -4 bp AA as the template (Fig. 3k). When the insertion size increased to 3 bp, the percentage of TIs decreased to 16.6% (Fig. 3l) but asymmetric TIs still existed (86.4% for -6/-5/-4 bp vs. 16.3% for -3/-2/-1 bp) (Fig. 3m). For instance, as shown at the target S33, 40.4% of all 3 bp insertions used -6 to -4 bp AGC as the template (Fig. 3n). As for TIs larger than 3 bp, the frequency was so rare that they were only detected at sporadic sites, as exemplified by a 5 bp TI at the S7 site (Fig. 3o). Altogether the above findings underscore the prevalence of TIs in CRISPR/Cas9 mediated insertions in MuSCs *in vivo*.

Lastly, we investigated potential factors associated with the formation of TIs. Overall, there was no clear correlation between the frequency of TIs and the TIF (Fig. 3p), indicating that the proportion of TIs is an inherent feature of each target site independent of the editing efficiency. While the frequency of total insertions did not show a significant association with local chromatin features (Fig. 3q), we found a striking negative correlation between the TI frequency and chromatin accessibility as measured by ATAC-seq (Fig. 3r). When exploring the relationship between TI formation and local DNA context, we found that similar to editing precision (Fig. 1t-1u), the base at the -3 bp position showed no significant association with TI frequency (Fig. 3s). However, sites with a G at the -4 bp position exhibited significantly lower TI frequency compared to those with A or T but not C (Fig. 3t). In sum, the above findings suggest that CRISPR/Cas9 mediated short insertions in MuSCs *in vivo* may be shaped by local DNA context at the DSB site.

### Limited predicting power of the machine learning models *in vivo*

Our above findings demonstrate that the main principles governing CRISPR/Cas9-mediated editing *in vitro* are largely applicable *in vivo*. This prompted us to test whether algorithms trained by profiling a large set of Cas9 target sites *in vitro* could be used for *in silico* prediction of the editing outcomes in MuSCs *in vivo*. To this end, we evaluated the performance of three widely used machine learning models—inDelphi^16^, FORECasT^17^, and Lindel^8^. We categorized all the detected indels into five groups based on types and size (> 2 bp deletions, 2 bp deletions, 1 bp deletions, 1 bp insertions and > 1 bp insertions) (Extended Data Fig. 4a). For each target site, the three algorithms were applied to predict the frequency of each indel group and compared with our experimental results. For > 2 bp deletions, 1 bp deletions, 1 bp insertions and > 1 bp insertions, inDelphi and FORECasT predicted significantly different frequencies compared to the experimentally measured; only for 2 bp deletions, yielded similar frequency as the measured (Extended Data Fig. 4a). Lindel performed slightly better, with its predicted frequencies for both the 2 bp deletions and the 1 bp insertions closely matching the experimental outcomes (Extended Data Fig. 4a). To further quantitatively assess the discrepancy between the predicted and measured indel frequencies, we computed the Pearson correlation coefficient for each site based on the five indel categories. All three models yielded moderate to strong correlations (0.64 for inDelphi, 0.67 for FORECasT, and 0.71 for Lindel) (Extended Data Fig. 4b), whereas FORECasT and Lindel have previously achieved an average Pearson coefficients exceeding 0.8 *in vitro*^12^. Collectively, these findings indicate that *in vitro*-trained algorithms have limited predictive power in forecasting indel patterns in MuSCs *in vivo*.

### *In vivo* mapping of CRISPR/Cas9 induced indel profiles in MuSCs using dual-sgRNAs

Next, we examined the Cas9 editing outcomes using dual sgRNAs. It is known that a single sgRNA-induced DSB can be repaired by NHEJ without editing, but Cas9 keeps cutting until an indel disrupts the target site^4,5,7^. Re-cutting can be prevented by employing two sgRNAs, as simultaneous editing can delete the intervening sequence so that the re-ligated sequence no longer aligns with the target sequences recognized by either sgRNA (Fig. 4a). In our previous studies, dual-sgRNA strategy has been frequently utilized to target a range of genomic regions to achieve successful deletion of various elements in MuSCs *in vivo*^31,33,34^. 51 pairs of sgRNAs were designed to target distinct genomic regions including those edited in our previous studies^31,33,34^, numerically labeled as D1 to D51 (Fig. 1a and Supplementary Table 1). Among them, 30 dual sgRNAs targeted CDS and promoter regions of 13 protein coding genes, including *Atf4*, *Bcl6*, *Ctcf*, *Ent*, *Fos*, *Fosb*, *Junb*, *Myc*, *Myod1*, *Pknox2*, *Pax7*, *Runx1* and *Sugt1*; 18 were designed to target potential enhancer regions, with three located near *Pax7* gene and four near *Runx1*; and 3 targeted non-regulatory intergenic regions to further increase the diversity and complexity of the targeted sites (Fig. 4b, Extended Data Fig. 5a and Supplementary Table 1). Considering the distance of the two concurrent DSBs may influence Cas9 editing outcomes^7^, the designed dual sgRNAs were separated by varying intervals mostly within 1 kb (Fig. 4c). Pax7^Cas9^ mouse at P10 was administrated with 5×10^11^ vg of the AAV-dual sgRNAs intramuscularly and MuSCs were isolated 4 weeks later (Fig. 1a). Sequences encompassing the two DSBs were amplified and subjected to unbiased NGS to detect the indel profiles (Supplementary Tables 2 and 5).

**Figure 4.**
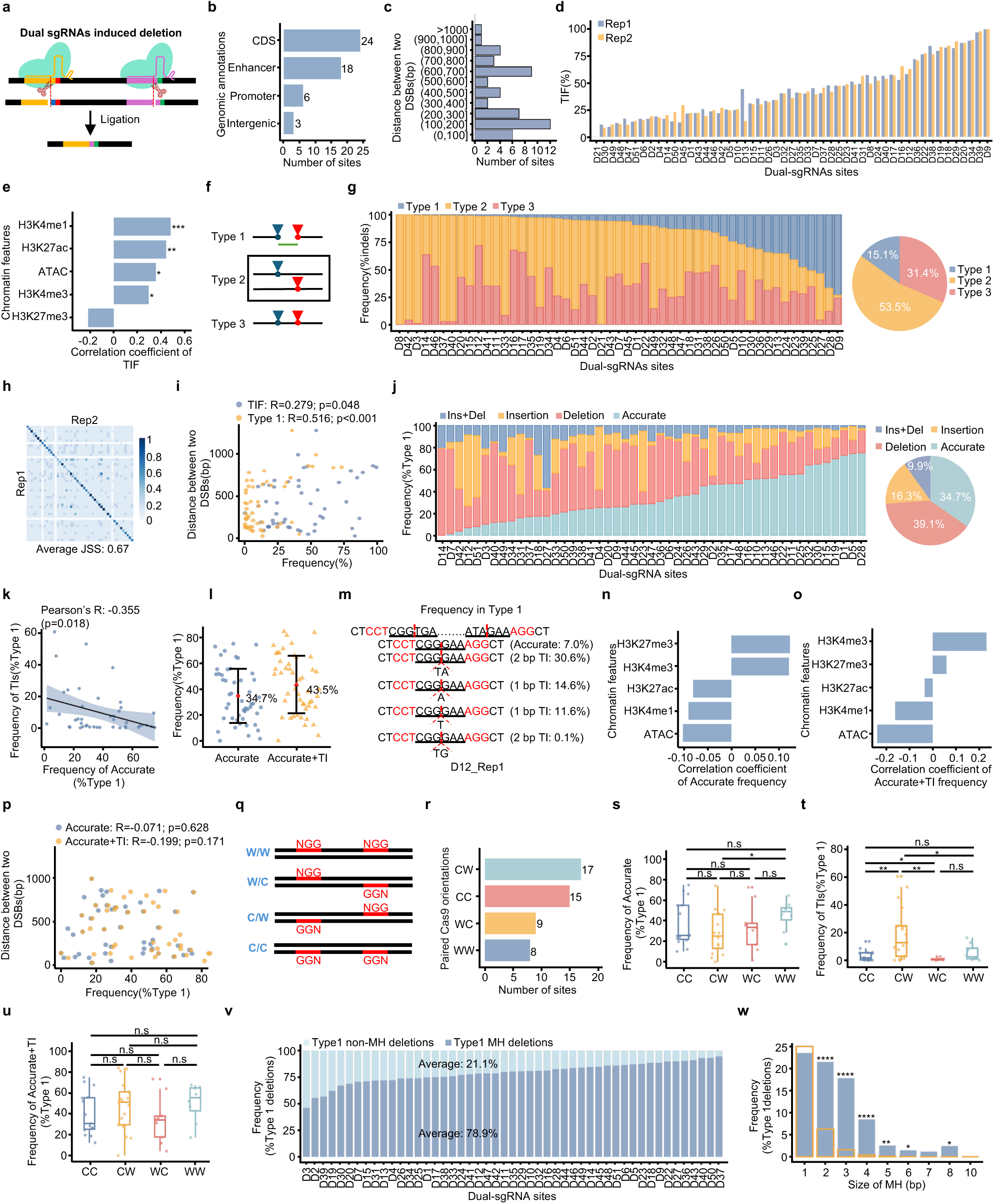
*In vivo* mapping of CRISPR/Cas9 induced indel profiles in MuSCs using dual-sgRNAs. (**a**) Schematic illustration of dual-sgRNAs induced DNA deletion. (**b**) The number of target sites with the indicated genomic annotations. (**c**) Distribution of distance between two DSBs across 51 sites. (**d**) TIFs for 51 dual-sgRNA targets with two biological replicates. (**e**) Spearman’s correlation between TIF and indicated chromatin features. (**f**) Schematic showing three types of dual-sgRNA induced indels. (**g**) Left: Frequency of Type 1, Type2 and Type 3 indels for each target. Right: Pie charting showing the average proportion of each type of indels across all targets. (**h**) Heatmap of JSSs for Type 1 indels. Each square indicates the JSS between the top five most frequent Type 1 indels of the corresponding targets. The average JSS of paired replicates across all targets is shown. (**i**) Correlation of distance between two DSBs and TIF (blue) or Type 1 (orange) frequency. (**j**) Left: Frequency of different subtypes of indel within Type 1 group for each target. Right: Pie chart showing the average frequency of different subtypes of indels in Type 1 group. 49 targets with over 100 Type 1 events were included for subtype analysis. (**k**) Pearson’s correlation between the frequency of TIs and Accurate indels in Type 1. Only 1 bp and 2 bp TIs were considered. (**l**) Frequency of Accurate alone and Accurate + TI in Type 1 indels. The average frequency across all targets is shown. (**m**) An example on D12 site showing the frequency of Accurate, 1 bp TI and 2 bp TI in Type 1. (**n-o**) Spearman’s correlation between the frequency of Accurate (**n**) or Accurate + TI (**o**) in Type 1 indels and indicated chromatin features. (**p**) Pearson’s correlation of distance between two DSBs and frequency of Accurate (blue) or Accurate + TI (orange). (**q**) Schematic showing four Cas9 PAM orientations for dual sgRNAs. W/W, W/C, C/W, and C/C orientations are determined by the recognition of Cas9 to the PAM motif on either the Watson strand (W) or the Crick strand (C). (**r**) Numbers of target sites categorized by each paired PAM orientation. (**s**-**u**) Boxplot illustrating the frequency of Accurate (**s**), TIs (**t**), Accurate + TI (**u**) in total Type 1 on targets with each PAM orientation. (**v**) Frequency of MH and non-MH mediated deletions in Type 1 deletions for each dual-sgRNA target. The average frequency of MH and non-MH mediated deletions across all sites is shown. (**w**) Frequency of deletions with MH of varying sizes in Type 1 deletions. The orange bar represents the expected frequency. Statistical significance was calculated using a t-test for (**s**) - (**u**) and a one-tailed t-test for (**w**). *p < 0.05, ** p < 0.01, ****p < 0.0001 and ns, no significance.

We found the dual-sgRNAs yielded a wide variation of TIFs ranging from 0.01% to 99.4% on the 51 target sites (Fig. 4d) with a much higher average of 42.5% compared to the single-sgRNA outcome (32.5%) (Fig. 1d), as noticed in our prior findings^31^. Similar to the findings from the single-sgRNA group (Fig. 1e), the TIFs positively correlated with active chromatin markers (Fig. 4e), further confirming that active chromatin may facilitate Cas9-mediated editing. Based on the occurrence of DSBs at the two target sites, we then categorized the repair outcomes into three types: 1) Both DSBs were formed simultaneously, resulting in the deletion of the intervening sequence; 2) Only one of the two sites was edited; 3) Editing occurs at both sites but DSBs were formed sequentially therefore no removal of the interval sequence (Fig. 4f). We found that Type 2 indels accounted for 53.5% of the total indels on average, followed by Type 3 of 31.4%, while Type 1 comprised only 15.1% (Fig. 4g) significantly lower than the 58.5% observed *in vitro*^7^. Similar to the single-sgRNA group (Extended Data Fig. 1c), the frequency of each type was comparable across the two replicates (Extended Data Fig. 5b). Type 1 indels resulted from ligation of the two DSBs to reflect a straightforward indication of Cas9 cleavage activity, thus becoming the focus of our subsequent analyses. Jaccard similarity scoring of the top five most frequent indels for each site—which represented an average of 62.9% of all Type 1 indels (Extended Data Fig. 6c-6d)—revealed an average similarity score as high as 0.67 between the two replicates (Fig. 4h and Supplementary Table 4); this indicates that Type 1 indel patterns were reproducible across replicates, reinforcing that Cas9 induced DNA repair in MuSCs is not random. Furthermore, the frequency of Type 1 indels did not show a significant correlation with chromatin features (Extended Data Fig. 5e). Nevertheless, we found that both TIFs and Type 1 frequency positively correlated with the distance between the two DSBs (Fig. 4i), supporting that the distance influences Cas9 induced repair outcomes^7^.

Next, to investigate the precision of NHEJ, we categorized the Type 1 indels into four subtypes, including Insertion, Deletion, Accurate (accurate ligation of the two broken ends) and Ins+Del (indels with simultaneous deletions and insertions) (Fig. 4j). Accurate indels reflect the precise nature of classical NHEJ^7^ and the frequency varied significantly across target sites, ranging from 0 to 75.1% (Fig. 4j), with an average of 34.7%; high similarity was detected across the two replicates (Extended Data Fig. 5f). It has been reported that the formation of TIs also reflects the precise nature of NHEJ and it suppresses the frequency of Accurate ligation events^7^; accordingly, we observed an inverse correlation between the frequency of the 1-2 bp TIs and the Accurate events (Fig. 4k). Therefore, the frequency of precise NHEJ-mediated repair reached 43.5% when considering Accurate and TIs together (Fig. 4l). As exemplified at the D12 site, both Accurate and TI existed and represented 7.0% and 56.9% of Type 1 indels, respectively (Fig. 4m). Of note, the average frequency of precise NHEJ *in vitro* can be 66.9%^7^, suggesting that Cas9-mediated editing *in vivo* is less precise.

Next, we explored potential factors influencing the NHEJ precision. Correlation analysis revealed that the frequency of Accurate or Accurate+TI indels exhibited no significant association with TIF (Extended Data Fig. 5g) or the frequency of Type 1 indels (Extended Data Fig. 5h), indicating that the proportion of precise NHEJ is an inherent property of each target site. Additionally, there were no correlations with chromatin features (Fig. 4n-4o) and the distance between the dual DSBs (Fig. 4p). Lastly, we investigated whether the orientation of dual PAM motifs might have any influence. The orientation determined by the recognition of Cas9 to the PAM motif on either the Watson strand (W) or the Crick strand (C), can be arranged in four distinct combinations: W/W, W/C, C/W, and C/C which were all included in our study (Fig. 4q-4r). W/C PAM configuration is shown to be more effective in generating accurate ligation than the other three^7^, which was however not observed in our study (Fig. 4s); Apart from a higher Accurate frequency associated with W/W compared to C/W, no significant differences were detected among the other PAM configurations (Fig. 4s). Interestingly, we found that the C/W orientation yielded the highest TI frequency and the W/C orientation produced the lowest (Fig. 4t), but no significant differences were detected in Accurate+TIs (Fig. 4u). These results suggest that PAM orientation configuration may not impact the NHEJ precision *in vivo*.

Additionally, we examined Type 1 deletions and found that on average 78.9% of Type 1 deletions exhibited MH (Fig. 4v), reinforcing that MH is widely used in deletion events. The length of the MH ranged from 1 to 10 bp, and the frequency of deletions was negatively correlated with the MH size but higher than the background expectation (Fig. 4w). Similar to the single-sgRNA group, the average size of Type 1 MH deletions was larger than that of Type 1 non-MH deletions (18 bp vs. 12 bp) (Extended Data Fig. 6a-6b). Furthermore, unsupervised hierarchical clustering revealed that Type 1 MH deletions were larger and peaked at 10-20 bp, with up to 63% exceeding 10 bp (Extended Data Fig. 6c). In contrast, Type 1 non-MH mediated deletions were generally smaller, with approximately 60% less than 10 bp (Extended Data Fig. 6d).

Altogether, the above results from analyzing dual sgRNAs mediated Cas9 editing outcomes demonstrate that the repair of Cas9-induced DSBs by NHEJ is inherently precise in MuSCs *in vivo* and also substantiate the extensive role of MH in Cas9-induced deletions.

### Analysis of CRISPR/Cas9 induced large on-target genomic modifications in MuSCs *in vivo*

Lastly, we focus our investigation on large-size genome modifications observed in our study (Fig. 1k). Emerging evidence demonstrates the presence of CRISPR/Cas9 induced rare large-size genomic modifications including LDs and LIs^13,19,22^, raising potential safety concerns in therapeutic genome editing. We first examined the LDs and LIs in the single sgRNA group and indeed LDs ≥ 50 bp occurred infrequently, accounting for an average of 3.2% of total indels and 4.5% of all deletion events (Fig. 5a) and the frequency was highly reproducible across the two biological replicates (Extended Data Fig. 7a). The number of distinct LD types varied substantially across target sites (Extended Data Fig. 7b). Jaccard similarity scoring of the LD types for each site revealed only modest similarity between the two replicates (average score: 0.31) (Fig. 5b and Supplementary Table 4), indicating that LD formation may be largely random. Consistent with previous study^10^, we found that 92.9% of LDs exhibited MH (Fig. 5c). Unlike typical MH mediated deletions, which predominantly involve 1 bp MH (Fig. 2c), these LDs preferentially utilized longer MH tracts (≥ 2 bp) (Fig. 5d); for example, the top three most frequent LD types (50, 54 and 69 bp) at S1 site were characterized by 3-4 bp MH (Extended Data Fig. 7c). Furthermore, LD frequency showed no significant association with TIF (Extended Data Fig. 7d), but positively correlated with active transcription with H3K4me3 signals (Fig. 5e). Given that longer indel size was correlated with lower editing precision and precise sites tended to produce short indels (Fig. 1r), we speculated that LD formation is associated with editing precision. As expected, the precise sites exhibited both a lower LD frequency and a reduced number of LD types (Fig. 5f) compared to the other two groups. Furthermore, the frequency of LD formation was inversely correlated with the frequency of the top 1 most frequent indel (Extended Data Fig. 7e), which was utilized to assess editing precision (Fig. 1o). Surprisingly, we found that the local DNA context surrounding the cut site also influenced LD formation: a G at the -3 bp position was typically associated with lower frequency of LDs compared to C (Fig. 5g, Left), whereas a -4 bp G correlated with elevated LD frequency compared to A and C (Fig. 5g, Right).

**Figure 5.**
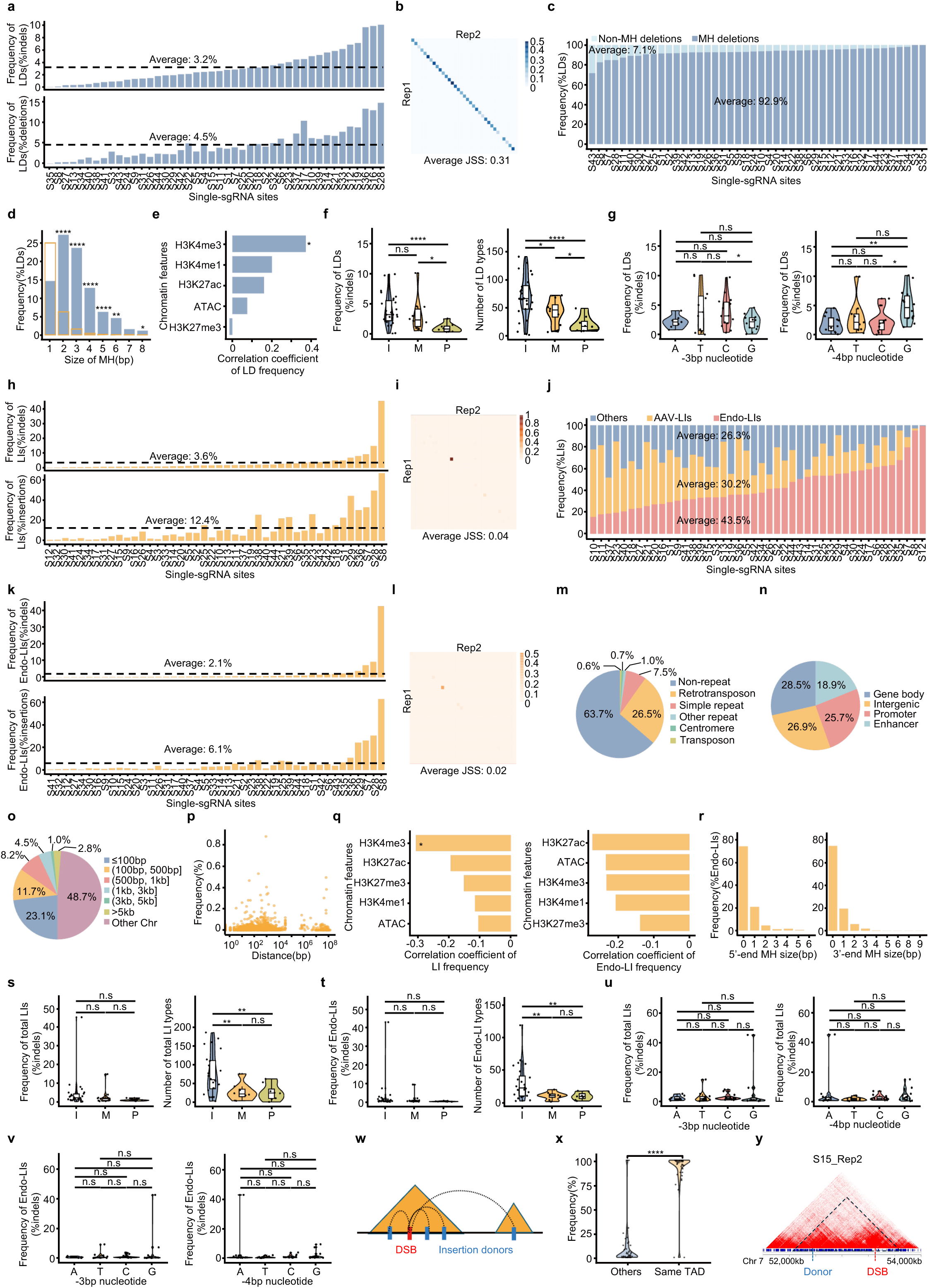
Analysis of CRISPR/Cas9 induced large on-target genomic modifications in MuSCs *in vivo*. (**a**) Frequency of ≥ 50 bp LDs for each single-sgRNA site, calculated as percentage in total indels (Upper) and total deletions (Lower). The average frequency of LDs across all sites is shown by the dash line. (**b**) Heatmap of JSSs for LDs. Each square shows the JSS between all LDs of corresponding targets. The average JSS of paired replicates across all targets is shown. (**c**) Frequency of MH and non-MH mediated LDs for each site. The average frequency across all sites is shown. (**d**) Frequency of LDs with MH of varying sizes. The orange bar represents the expected frequency for LDs with each MH size. (**e**) Spearman’s correlation between LD frequency in total indels and indicated chromatin features. (**f**) Violin plot illustrating the frequency of LDs (Left) and number of LD types (Right) in imprecise (I), middle (M), precise (P) groups. (**g**) Frequency of LDs in total indels at sites with A, T, C, G at -3 bp (Left) of -4 bp (Right) upstream of the PAM motif. (**h**) Frequency of ≥ 20 bp LIs for each single-sgRNA site, calculated as percentage in total indels (Upper) and total insertions (Lower). The average frequency of LIs across all sites is shown by the dash line. (**i**) Heatmap of JSSs for LIs. Each square shows the JSS between all LIs of corresponding targets. The average JSS of paired replicates across all targets is shown. (**j**) Frequency of different types of LIs. The average frequency of each type across all sites is shown. (**k**) Frequency of ≥ 20 bp Endo-LIs for each single-sgRNA site, calculated as percentage in total indels (Upper) and total insertions (Lower). The average frequency is shown by dash line. (**l**) Heatmap of JSSs for Endo-LIs. Each square shows the JSS between all Endo-LIs of corresponding targets. The average JSS of paired replicates across all targets is shown. (**m**) Pie chart showing the frequency of donor sequence categorized by types of repetitive elements for Endo-LIs. (**n**) Pie chart showing the frequency of donor sequence categorized by genomic annotations for Endo-LIs. (**o**) Pie chart showing distribution of distances between insertion donors for Endo-LIs and associated DSBs. (**p**) Scatter plot illustrating the distance distribution between donor sequences for Endo-LIs and their associated DSBs on the same chromosome. (**q**) Spearman’s correlation between the frequency of total LIs (Left) or Endo-LIs (Right) and indicated chromatin features. (**r**) Frequency of MH with varying sizes in 5’ (Left) and 3’ (Right) ends of Endo-LIs. (**s**) Violin plot illustrating the frequency of total LIs (Left) and number of total LI types (Right) in imprecise (I), middle (M), precise (P) groups. (**t**) Violin plot illustrating the frequency of Endo-LIs (Left) and number of Endo-LI types (Right) in imprecise (I), middle (M), precise (P) groups. (**u**) Frequency of total LIs at sites with A, T, C, G at -3 bp (Left) of -4 bp (Right) upstream of the PAM motif. (**v**) Frequency of Endo-LIs at sites with A, T, C, G at -3 bp (Left) of -4 bp (Right) upstream of the PAM motif. (**w**) Schematic showing co-localization of donor sequences and associated DSBs of Endo-LIs within the same TAD. (**x**) Violin plot showing significantly higher Endo-LI frequency of donor sequences originating from DSB-containing TADs compared to other loci on the same chromosome. (**y**) An example on S15 illustrating the influence of TADs on Endo-LI formation. The positions of the donor and DSB are indicated. The dashed triangle denotes the identified TAD. Statistical significance was calculated using a t-test for (**f**), (**g**), (**s**) - (**v**) and a paired t-test for (**x**) and a one-tailed t-test for (**d**). *p < 0.05, **p < 0.01, ****p < 0.0001 and ns, no significance.

Similar to LDs, LIs ≥ 20 bp occurred rarely, representing only 3.6% of total indels and 12.4% of all insertion events (Fig. 5h) and showed high reproducibility across replicates (Extended Data Fig. 7f); Also, the number of LIs varied substantially across all sites (Extended Data Fig. 7g). Unexpectedly, a minimal similarity in LI types was observed between the two replicates (average JSS: 0.04) (Fig. 5i and Supplementary Table 4), suggesting that the formation of LIs may be stochastic and highly complex. Further analysis of the origins of the inserted sequences revealed that the majority (73.7%) of LIs were templated, with 43.5% originating from the mouse genome (endogenous LIs, Endo-LIs) and 30.2% from the AAV vector (AAV-LIs) (Fig. 5j). Integration from viral vectors has been extensively studied^13,19^, while Endo-LIs derived from the host genome remain understudied. Overall, the Endo-LIs accounted for 2.1% of the total indels and 6.1% of all insertions (Fig. 5k); they exhibited characteristics similar to the total LIs, including a reproducible frequency between the two replicates (Extended Data Fig. 7h), variation in the number of types across sites (Extended Data Fig. 7i), and a lack of similarity in Endo-LI patterns (Fig. 5l and Supplementary Table 4). Notably, a fraction of Endo-LIs originated from repetitive sequences and fragile genomic regions, such as retrotransposons (26.5%), centromeres (0.7%) (Fig. 5m). Recent reports have suggested that DSB-induced insertions are associated with active transcription^26^. Consistently, 28.5% and 26.9% of the detected donor sequences came from promoters and gene body regions, respectively (Fig. 5n). Moreover, we found that a subset of Endo-LIs (18.9%) originated from enhancer regions (Fig. 5n), suggesting a possibility for them to modulate nearby gene expression. In general, over 98% of Endo-LIs originated from a single donor locus, while a small portion derived from up to four genomic loci (Extended Data Fig. 7j). Regarding the distance from DSBs, 48.7% of Endo-LIs originated from different chromosomes (Fig. 5o and Extended Data Fig. 7k), and the other half were located on the same chromosome, particularly within 5 kb of the DSB (Fig. 5o-5p and Extended Data Fig. 7k).

Furthermore, no significant correlation was observed between the frequency of total LIs or Endo-LIs with TIF (Extended Data Fig. 7l). In contrast to LDs (Fig. 5e), the total LI frequency exhibited a negative correlation with active transcription marked by H3K4me3 (Fig. 5q, Left), although Endo-LIs showed no significant bias (Fig. 5q, Right). Further investigation of MH involvement revealed that about 74% of Endo-LIs showed no detectable MH at either 5′ or 3′ ends (Fig. 5r), indicating MH was not prevalently utilized in Endo-LIs formation and MMEJ may not be the predominant repair pathway for Endo-LIs. Moreover, there was no clear association between the formation of both total and Endo-LIs with editing precision as comparable frequency of total and Endo-LIs were detected at I, M and P sites (Fig. 5s-5t and Extended Data Fig. 7m-7n) although I sites generated higher numbers of total LI and Endo-LI types (Fig. 5s-5t). Additionally, the flanking sequence composition at the cut site had no effect on total and Endo-LI frequency (Fig. 5u-5v). It has been reported that insertions require physical contact between donor and acceptor loci^26^ and a recent study also showed that Topologically Associated Domains (TADs) can restrict DSB induced DNA replication inhibition^44^, we thus examined the possible impact of TAD organization on Endo-LI formation (Fig. 5w). By analyzing the Hi-C data from primary myoblasts^42^, we found that a significantly higher frequency (90.5% vs. 9.5%) of donor sequences and their associated DSBs originated from the same TAD (Fig. 5x and Extended Data Fig. 10o) as exemplified on S15 in Fig. 5y, suggesting TAD organization may play a role in regulating Endo-LIs formation.

Next, we examined the formation of large genome modifications in the dual-sgRNA group (Extended Data Fig. 8) and found that LDs accounted for an average of 2.5% of Type 1 indels and 3.5% of Type 1 deletion events (Extended Data Fig. 8a). Also, MH was prevalent during LD formation, with 87.5% of LDs exhibiting MH (Extended Data Fig. 8b), and these LDs preferentially utilized longer MH tracts (≥ 2 bp) (Extended Data Fig. 8c), further underscoring the role of MMEJ in their generation. LIs constituted 1.0% of total indels and 5.9% of all insertion events (Extended Data Fig. 8d). 28.7% of LIs originated from the mouse genome, while 41.9% were derived from the AAV viral vector (Extended Data Fig. 8e). 38.1% of donor sequences of Endo-LIs were from fragile genomic regions, including retrotransposons (22.8%), centromeres (1.0%) (Extended Data Fig. 8f), and 10.88% from enhancer regions (Extended Data Fig. 8g). Of note, the role of the 3D genome in Endo-LIs formation was further supported by the observation that donor sequences and associated DSBs largely co-localize in the same TAD (Extended Data Fig. 8h-8i).

Altogether, the above results demonstrate that CRISPR/Cas9-induced large on-target genomic modifications can occur in MuSCs *in vivo*. LD formation requires the presence of MH and is influenced by the local DNA context, while the formation of Endo-LIs is affected by 3D chromatin architecture.

### AAV integration is a general outcome of CRISPR/Cas9-induced editing in MuSCs *in vivo*

We next focused our attention on exogenous AAV-LIs which is emerging as a common phenomenon in AAV-delivered CRISPR/Cas9 editing^9,29,30,43^. AAV-LIs were detected in 43 out of the 44 sites, accounting for an average of 0.9% of total indels in single-sgRNA group (Fig. 6a). We suspect this may be underestimated as only integrations ≥ 20 bp were used in our analysis. Similarly, the frequency of AAV-LI was highly consistent across the two replicates (Extended Data Fig. 9a). Moreover, insertion positions at both the 5′ and 3′ ends were predominantly located adjacent to the CRISPR/Cas9 cut site (Fig. 6b), indicating these integrations were indeed generated by CRISPR/Cas9-induced DNA breaks.

**Figure 6.**
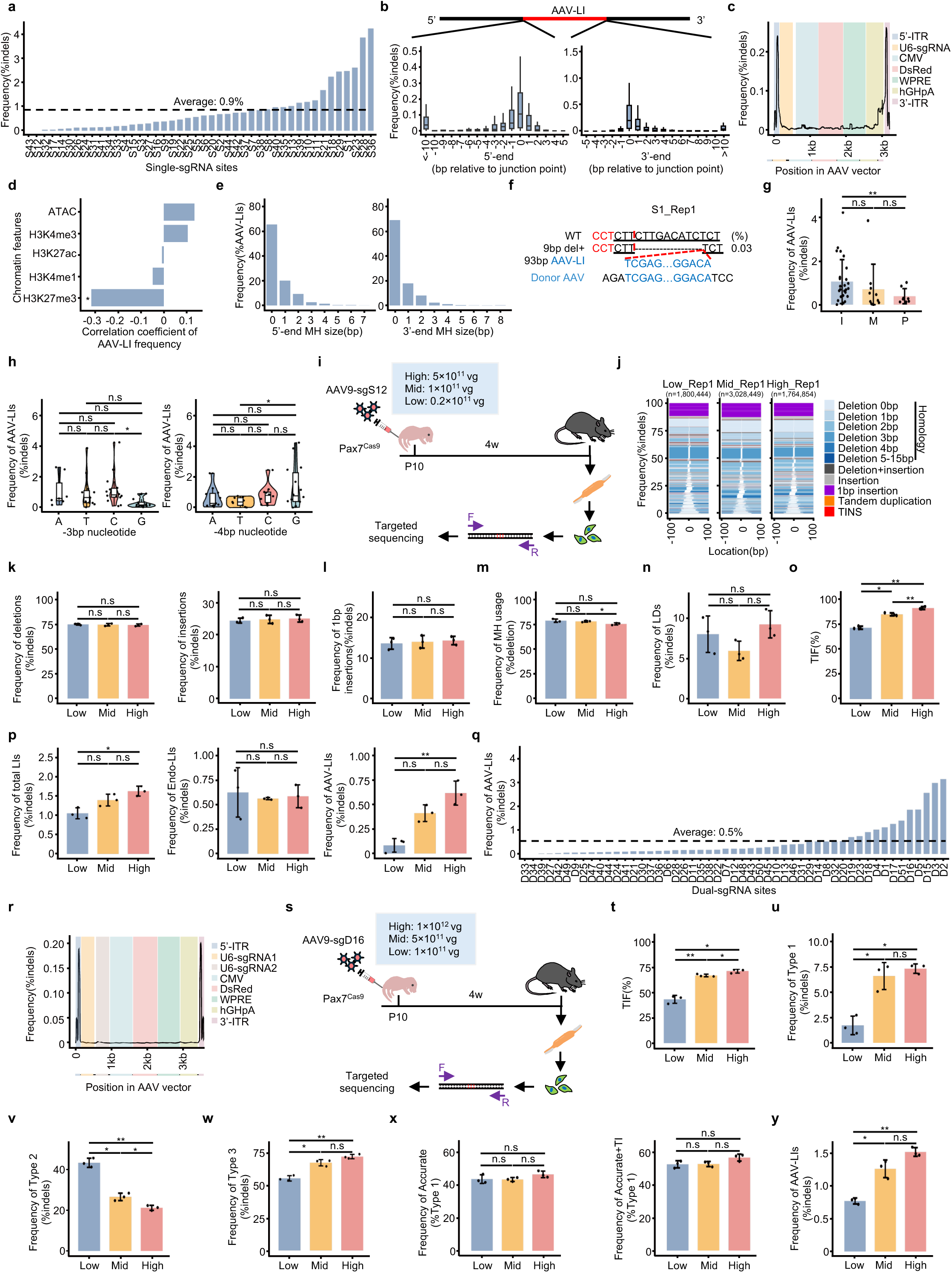
AAV integration is a general outcome of CRISPR/Cas9-induced editing in MuSCs *in vivo*. (**a**) Frequency of AAV-LI for each single-sgRNA site, calculated as percentage in total indels. The average frequency of AAV-LIs across all sites is shown by the dash line. (**b**) Distribution of 5’ and 3’ end of AAV integration sites relative to the junction point. (**c**) The average inserted frequency of each AAV vector region. Each element of the AAV vector is presented by designated color and the position in the AAV vector is shown. (**d**) Spearman’s correlation between AAV-LI frequency and the indicated chromatin features. (**e**) Frequency of MH with varying sizes in 5’ (Left) and 3’ (Right) ends of AAV-LI. (**f**) An example of AAV-LI on S1. A 93 bp AAV integration occurred together with a 9 bp deletion at the cut site. Frequency is presented as a percentage in total indels. The AAV-LI sequence and donor AAV sequence is marked in blue. (**g**) Violin plot illustrating the frequency of AAV-LI in I, M and P groups. (**h**) Frequency of AAV-LI at sites with A, T, C, G at -3 bp (Left) or -4 bp (Right) upstream of the PAM motif. (**i**) Schematic showing experimental design for assessing the effect of AAV dosage on AAV-LI at the S12 site. High (5×10^11^ vg), Mid (1×10^11^ vg) or Low (0.2×10^11^ vg) dose of AAV9-sgRNA virus containing sgRNA targeting the S12 site were IM injected into Pax7^Cas9^ mouse at P10, with three replicates for each, followed by MuSC isolation four weeks after injection. Sequences encompassing the S12 site were amplified and subjected to unbiased deep-sequencing. (**j**) Visualized plots illustrating the indel profiles for Rep_1 of S12 target in each AAV dosage group. (**k**-**p**) Barplot illustrating frequency of deletions (**k**, Left), insertions (**k**, Right), 1 bp insertions (**l**), MH usage in deletions (**m**), LDs (**n**), TIFs (**o**), total LIs (**p**, Left), Endo-LIs (**p**, Middle) and AAV-LIs (**p**, Right) for the three dosage groups (n=3 mice in each group). (**q**) Frequency of AAV-LIs in dual-sgRNA group, calculated as percentage in total indels. The average frequency of AAV-LIs across all sites is shown by the dash line. (**r**) Average insertion frequency of indicated AAV–dual sgRNA vector regions. (**s**) Schematic showing experimental design for assessing the effect of AAV dosage on AAV-LI at the D16 site. High (1×10^12^ vg), Mid (5×10^11^ vg) or Low (1×10^11^ vg) dose of AAV9-dual sgRNA virus targeting the D16 site were IM injected into Pax7^Cas9^ mouse at P10, (n=3 mice), followed by MuSC isolation four weeks after injection. Sequences encompassing the D16 site were amplified and subjected to unbiased deep-sequencing and analysis. (**t**-**y**) Barplot illustrating TIF (**t**), frequency of Type 1 indels (**u**), Type 2 indels (**v**), Type 3 indels (**w**), Accurate events (**x**, Left), Accurate + TI events (**x**, Right) and AAV-LIs (**y**) for the three dosage groups (n=3 mice in each group). Statistical significance was calculated using a t-test for (**g**) - (**h**) and a paired t-test for (**k**) - (**p**) and (**t**) - (**y**). *p < 0.05, **p < 0.01 and ns, no significance.

Previous studies have reported that the inverted terminal repeat (ITR) region of AAV is the hotspot for vector integration^29^. Accordingly, we found that the ITR elements at both ends of the AAV vector were more frequently enriched in AAV-LI reads compared to other vector elements such as the U6-sgRNA cassette, CMV-DsRed cassette, WPRE and hGHpA (Fig. 6c). Similar to total LIs, the frequency of AAV-LIs showed no significant correlation with TIF (Extended Data Fig. 9b) and was negatively correlated with H3K27me3 but not other chromatin signals (Fig. 6d). Regarding the role of MH in AAV-LIs, we observed that the majority (> 65%) of AAV-LIs contained no MH at 5’ or 3’ end (Fig. 6e), indicating MH may not be essential for AAV integration; this was exemplified at the S1 site, a 93 bp fragment of the AAV vector was integrated at the cut site accompanied by a 9 bp deletion but no MH detected at either end of the AAV-LI (Fig. 6f). Additionally, the frequency of AAV-LIs was negatively correlated with editing precision, with precise sites exhibiting significantly lower average frequency in comparison to the imprecise group (Fig. 6g). The sequence context flanking the cut site also influenced AAV-LIs: a G at the –3 bp position was associated with reduced integration compared to C (Fig. 6h, Left), whereas a G at the –4 bp position showed enhanced AAV-LI frequency compared to T (Fig. 6h, Right).

As AAV dosage is a crucial consideration in AAV mediated gene therapy^28^, we examined the AAV dosage effect on AAV integration leveraging our previously collected sequencing data performed on the single sgRNA S12 site (located in the CDS region of *Myod* locus) using low (0.2×10^11^ vg), medium (1×10^11^ vg), or high (5×10^11^ vg) AAV doses^31^ (Fig. 6i). Reanalysis of the data (Supplementary Tables 2 and 6) revealed that although the overall indel profile— including the general editing patterns (Fig. 6j), frequency of deletions (Fig. 6k, Left), insertions (Fig. 6k, Right), 1 bp insertions (Fig. 6l), MH usage in deletions (Fig. 6m) as well as LDs (Fig. 6n)—was largely unaffected, increasing the AAV dose indeed enhanced the TIF (Fig. 6o). As expected, the overall frequency of total LIs was significantly higher in the high-dose compared to the low-dose group (Fig. 6p, Left), primarily driven by elevated AAV integration (Fig. 6p, Right), whereas the frequency of Endo-LIs remained largely unchanged (Fig. 6p, Middle). The above findings suggest that while increasing AAV dosage can enhance editing efficiency in AAV-mediated CRISPR/Cas9 *in vivo* editing, it may introduce unwanted risks by elevating the frequency of AAV integration.

Dual-sgRNA group was also examined by aligning AAV sequences with sequencing reads. We found that AAV integration occurred at all 51 sites, with an average frequency of 0.5% of total indels (Fig. 5q) and comparable across replicates (Extended Data Fig. 9c). Consistent with the single-sgRNA group (Fig. 6c), most of the mapping reads included the ITR element (Fig. 6r). Furthermore, increasing AAV dosage was associated with higher TIF frequency (Fig. 6s-6t, Supplementary Tables 2 and 6) and led to a significant increase in the frequency of Type 1 and Type 3 but not Type 2 events (Fig. 6u-6w). We reason that the formation of Type 1 and Type 3 requires the combined action of two sgRNAs which may be facilitated by the increased AAV-sgRNA dose. However, the dosage did not affect the frequency of accurate NHEJ events within the Type 1 indels as the levels of Accurate and Accurate+TI were comparable among the three dosage groups (Fig. 6x). As expected, increasing the dosage from low to high elevated AAV-LI formation (Fig. 6y).

In summary, our findings demonstrate that AAV integration is a commonly detected outcome of CRISPR/Cas9-induced editing in MuSCs *in vivo*, highlighting the need to take caution when using AAV-mediated CRISPR/Cas9 therapy *in vivo*.

### *In vivo* perturbation of NHEJ/MMEJ pathways can modulate CRISPR/Cas9 editing in MuSCs

Given that NHEJ and MMEJ pathways dominate repair outcomes in the absence of exogenous donor templates during *in vivo* genome editing, we next investigated their contributions to large on-target modifications by depleting key components of each pathway (Fig. 7a). To this end, as shown in Fig. 7b, AAV9-dual sgRNA (8×10^11^ vg) targeting key factors of NHEJ (Ku80, Lig4) or MMEJ (Nbn, Polq) were intramuscularly injected into Pax7^Cas9^ mice at P5; AAV9-single sgRNA (5×10^11^ vg) was administrated at P10 to target the above studied S29 site which exhibited moderate levels of LDs (Fig. 5a) and LIs (Fig. 5h, 5k and 6a). MuSCs were isolated four weeks later and near depletion of the above proteins in MuSCs was confirmed (Fig. 7c).

**Figure 7.**
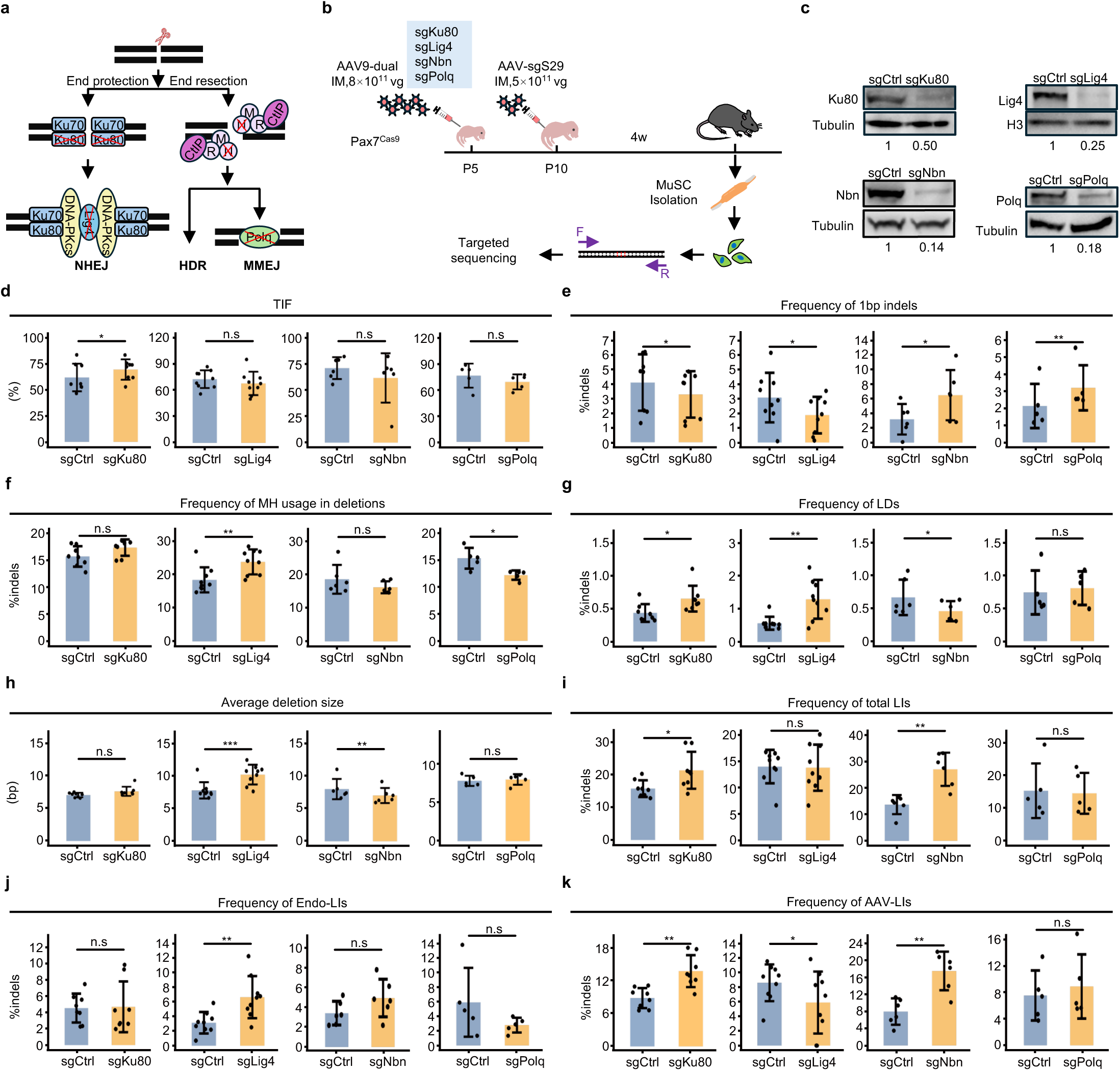
*In vivo* perturbation of NHEJ/MMEJ pathways can modulate CRISPR/Cas9 editing in MuSCs. (**a**) Schematic illustration of CRISPR/Cas9 induced DSB repair pathways. Key proteins in each pathway are shown and proteins selected for *in vivo* deletion are marked in red. (**b**) Schematic showing experimental design for deleting the above proteins in MuSCs *in vivo*. 8×10^11^ vg of AAV9-dual sgRNA virus targeting the CDS regions for Ku80, Lig4, Nbn, or Polq were IM injected into Pax7^Cas9^ mouse at P5, respectively. 5×10^11^ vg of AAV9-single sgRNA was administrated at P10 to target the S29 site. MuSCs were isolated four weeks after injection and sequences encompassing the S29 site were amplified and subjected to unbiased deep-sequencing. (**c**) Protein depletion of Ku80, Lig4, Nbn, or Polq in the above MuSCs was confirmed by Western blotting. (**d**-**k**) TIFs (**d**), frequency of 1 bp indels (**e**), MH usage (MH size ≥ 2) in deletions (**f**), LDs (**g**), total LIs (**i**), Endo-LIs (**j**), AAV-LIs (**k**) and average deletion size (**h**) after each protein depletion is shown. Statistical significance was calculated using a paired t-test for (**d**) - (**k**) (n=5 to 9 in each group). *p < 0.05, **p < 0.01, ***p < 0.001 and ns, no significance.

Targeted sequencing was performed to examine how each perturbation altered editing outcomes (Supplementary Tables 2 and 7). Notably, only the loss of Ku80 but not others significantly increased the TIF (Fig. 7d). Both Ku80 and Nbn depletion, but not Lig4 or Polq, led to a marked reduction in the frequency of deletions among total indels (Extended Data Fig. 10a). As expected, 1 bp indels—typically a hallmark of NHEJ—were significantly decreased following Ku80 or Lig4 depletion but increased upon Nbn or Polq loss (Fig. 7e). Conversely, Loss of Lig4 enhanced the average usage MH in deletions, hinting elevated MMEJ activity (Fig. 7f); but Polq depletion had the opposite effect; Ku80 or Nbn loss also led to expected decrease and increase respectively although without statistical significance (Fig. 7f). These findings suggest that disrupting these factors *in vivo* can modulate the activity of their respective repair pathway.

We next assessed how the perturbations influenced the formation of large on-target DNA modifications. Disruption of NHEJ components Ku80 or Lig4 significantly increased the frequency of LDs; Nbn depletion decreased LD occurrence but Polq loss had minimal impact despite its established role in promoting LDs *in vitro*^10^(Fig. 7g). Additionally, loss of Lig4 increased the average deletion size, but Nbn depletion caused opposite effect (Fig. 7h). The effects on LIs were complex (Fig. 7i-7k): Ku80 and Nbn loss elevated both the frequency of total LIs (Fig. 7i) and AAV-LIs (Fig. 7k), without affecting the proportion of Endo-LIs (Fig. 7j). Polq depletion had a minimal impact on total LIs, Endo-LIs, or AAV-LIs (Fig. 7i-7k). Interestingly, Removal of Lig4 did not alter total LIs (Fig. 7i) but significantly increased Endo-LIs (Fig. 7j) and decreased AAV-LIs (Fig. 7k). Moreover, none of the perturbations affected the overall insertion size or the size of endogenous insertions (Extended Data Fig. 10b-10c). Collectively, our results demonstrate that dampening NHEJ activity promotes the LD formation, whereas suppression of MMEJ causes opposite effect. Furthermore, distinct NHEJ and MMEJ factors differentially influence LI generation, suggesting a more complex mechanistic basis for LI formation *in vivo*.

## Discussion

In this study, by systematically investigating the editing patterns at 95 sites in MuSCs *in vivo*, we revealed the general principles of CRISPR/Cas9-mediated *in vivo* editing. Our results indicate that the repair outcomes of CRISPR/Cas9 generated DSB in MuSCs *in vivo* largely align with those observed *in vitro*. Large on-target genome modifications are widely present, representing an important class of editing byproducts that may pose potential risks for CRISPR/Cas9-mediated therapies. Furthermore, through depleting key factors in NHEJ and MMEJ pathways, we uncovered their distinct roles in the generation of these undesirable byproducts.

### *In vivo* CRISPR/Cas9 editing in MuSCs follows *in vitro* principles but with reduced precision

Although the main rules of CRISPR/Cas9-induced indel signatures derived from *in vitro* studies generally persist *in vivo*—such as the non-random nature of indel profiles (Fig. 1g), the prevalence of MH in deletions (Fig. 2b and 4v), and the favor of NHEJ for precise editing (Fig. 3 and Fig. 4)—there are notable differences between *in vitro* and *in vivo* editing. First, *in vivo* editing appears to be less precise compared to *in vitro* editing; this is evidenced by the largely increased number of sites with imprecise editing in the single-RNA group (Fig. 1o) and reduced frequency of accurate NHEJ in dual-sgRNA group (Fig. 4l). Second, the directionality of deletions *in vivo* is distinct. Previous *in vitro* study using massively parallel synthetic reporters containing a sgRNA expression cassette together with its PAM-endowed target sequence demonstrates that both MH- and non-MH-mediated deletions exhibit asymmetric characteristics, predominantly occurring unidirectionally downstream of the breakpoint^8^, but we observed that the overall deletions (Fig. 2j) and MH mediated deletions (Fig. 2k) preferred to span the cut sites. In contrast, non-MH mediated deletions show a higher tendency for occurring upstream (Fig. 2l). Furthermore, when we applied the machine learning models trained on *in vitro* datasets to predict the *in vivo* indel patterns, the performance was suboptimal (Extended Data Fig. 4). Our findings thus demonstrate that the *in vivo* repair of CRISPR/Cas9-induced DSBs is inherently more complex; principles extracted from the *in vitro* studies using synthetic reporters may not extrapolate to assess editing in a native epigenetic environment or heterogeneous *in vivo* microenvironment.

### NHEJ-mediated CRISPR/Cas9-induced DSB repair *in vivo* is precise and sequence context dependent

Our findings uncover that deletions predominate the landscape of CRISPR/Cas9-induced indels in MuSCs (Fig. 1f). Both MMEJ and NHEJ can lead to deletions, and NHEJ is thought to be primarily responsible for generating small deletions^5^, typically within 2 bp, whereas larger size deletions are mainly formed by MH mediated MMEJ. Most deletions in our analysis are accompanied by MH (Fig. 2b and 4v), implying MMEJ could be the predominant pathway for these deletions. Despite high reproducibility across replicates (Fig. 1g), increased deletion rates potentially compromise editing precision to some extent (Fig. 1s). In contrast, NHEJ-mediated small insertions in our study tended to be accurate, often manifested as TIs (Fig. 3 and Fig. 4). In CRISPR/Cas9-related *ex vivo* gene therapies, enhancing HDR levels by simultaneously inhibiting MMEJ and NHEJ is a commonly used strategy to increase the editing precision^5^. *In vivo* CRISPR/Cas9 based therapies on the other hand primarily rely on the NHEJ and MMEJ pathways since HDR is largely inactive in terminally differentiated cells or requires an exogenous repair template^9^. Compared to MMEJ, NHEJ tends to produce smaller and more predictable outcomes, making it a safer option for *in vivo* editing; thus, strategies to enhance accurate NHEJ may be beneficial for *in vivo* therapies. For example, the Plk3 inhibitor GW843682X can enhance accurate NHEJ by repressing CtIP-dependent end resection^7^.

On the other hand, due to the cleavage flexibility of the RuvC domain, Cas9 can produce both blunt and staggered ends^3,40^. Previous studies have shown that blunt ends, upon repair, may result in random deletions, template-independent insertions, or restoration of the wild-type sequence^3^. In contrast, the generation of an overhang typically results in TIs through NHEJ repair. Emerging studies have further demonstrated that the type of end generated is closely related to the DNA sequence context near the breakpoint^3,40^. Increasing evidence suggests that when the nucleotide at the -4 bp upstream of the PAM motif is G, the RuvC domain is more inclined to cleave the non-target DNA strand to produce blunt DSBs, whereas a G at -3 bp is more frequently linked to staggered DSBs^15,16,40^. Consistent with these observations, our results show that target sites harboring a −3 bp G were generally associated with enhanced editing precision (Fig. 1t), reduced frequency of LDs (Fig. 5g, Left) and AAV integration (Fig. 6h, Left). In contrast, a −4 bp G displayed the opposite pattern, characterized by reduced editing precision (Fig. 1u), decreased proportion of TIs (Fig. 3t), and elevated frequency of both LDs (Fig. 5g, Right) and AAV integration (Fig. 6h, Right). Therefore, during *in vivo* editing, alongside enhancing NHEJ activity, selecting target sites with specific sequence features could increase the generation of staggered DSBs, further improving editing precision.

### Widespread occurrence of large on-target genome modifications in CRISPR/Cas9 mediated *in vivo* editing in MuSCs

Our study reveals widespread occurrence of large on-target modifications during *in vivo* editing. With the advent of long-read sequencing, LDs and LIs, including the integration of exogenous vector DNA, have been progressively reported^10,13,23^ but mostly on sporadic sites *in vitro*. Our work demonstrates that these large on-target modifications are also prevalent during *in vivo* editing. Among these modifications, LDs are the most extensively studied. Their occurrence can result in the loss of large DNA fragments or even the removal of entire chromosomes in some cases^21,22^. In 92.9% of the LDs analyzed (Fig. 5c), we observed the presence of MH, implicating the involvement of MMEJ pathway in their formation, which aligns with recent studies^10,13^. Of note, the indels were detected four weeks after sgRNA transduction in MuSCs, suggesting that these undesirable mutations can persist stably.

Current studies on LIs were primarily focused on the integration of exogenous DNA vectors, such as AAV^13,29,30^. As one of the most widely utilized delivery platforms in clinical gene therapy, AAV integration at target sites has been reported during *in vivo* editing at sporadic sites, with integration rates at certain loci reaching as high as 47% of all indels^29^. Our results indicated that AAV integration is also a common phenomenon across the wide range of sites during *in vivo* editing in MuSCs (Fig. 6a and 6q), with the ITR region identified as a hotspot for integration (Fig. 6c and 6r). Notably, the AAV integration accounted for only 0.9% of all indels on average, with the highest only 4.2% (Fig. 6a). Several factors may contribute to this discrepancy from the *in vitro* finding, including differences in cell types, AAV dosage, and most notably, analytical criteria. In our analysis, we only considered AAV integrations greater than 20 bp as a subset of LIs, which may cause the underestimation. Our findings thus reinforce the notion that the potential safety concerns of AAV integration remain to be thoroughly evaluated; the use of non-viral vectors should be considered for the transfer of gene editing agents *in vivo*, such as lipid nanoparticles (LNPs) for Cas9 mRNA delivery^28^.

Previous studies have shown that a portion of LIs can also originate from the host genome^13,23^, but there remains a significant knowledge gap in understanding these Endo-LIs. Among the LIs detected in the single-sgRNA group, 43.5% of the donor sequences were derived from the host genome (Fig. 5j). Notably, a subset of Endo-LIs sourced from enhancer regions (Fig. 5n and Extended Data Fig. 8g), suggesting their potential role in disrupting gene regulation. For example, if DSBs occur near oncogenes, such insertions could enhance oncogene expression, potentially contributing to tumorigenesis. Further analysis of these Endo-LI types revealed no reproducibility across biological replicates (Fig. 5l), indicating a high degree of randomness or complexity in their generation. Nevertheless, we found that the 3D genome architecture influenced the formation of these insertions. Specifically, insertions originating from the same chromosome were more likely to reside within the same TAD as the acceptor site (Fig. 5x and Extended Data Fig. 8i). This observation aligns with the expectation that endogenous insertion events typically require physical proximity between donor and acceptor loci^26^; it is also consistent with a recent study showing that TAD can restrict DSB induced DNA replication inhibition^44^. Future studies are necessary to elucidate the mechanisms underlying the formation of these Endo-LIs and assess their potential safety risks.

### Differential roles of NHEJ/MMEJ in generating large on-target genome modifications in MuSCs *in vivo*

Despite the increasing attention on Cas9 generated large gene modifications^13,20,23^, the mechanisms underlying their formation remain elusive. Here we depleted four key factors essential for NHEJ and MMEJ to investigate their involvement. Our findings indicate that depleting Ku80 or Lig4 to reduce NHEJ activity resulted in an increased LD formation (Fig. 7g). Similarly, depleting the key protein Nbn involved in MMEJ resulted in a decreased LD frequency (Fig. 7g). Interestingly, while *in vitro* studies have shown that the knockout or inhibition of Polq significantly reduces LD formation^10^, we did not observe such phenomenon *in vivo* (Fig. 7g), although we noted a reduction in MH-mediated deletions following Polq loss (Fig. 7f). This discrepancy might arise from incomplete knockout of Polq (Fig. 7c) or differences in DNA repair pathways between the *in vivo* and *in vitro* contexts. In terms of LIs, their formation appears to be more complex. The loss of Ku80 and Nbn both resulted in an increased frequency of total LIs (Fig. 7i), likely due to elevated AAV integration (Fig. 7k), but the formation of Endo-LIs remained unchanged (Fig. 7j). Of note, since an AAV-dual sgRNA strategy was used to knock out these factors, we cannot rule out the possibility that variation in AAV transduction efficiency among mice caused changes in AAV integration. While studies have indicated that both MMEJ and NHEJ are involved in the random integration of exogenous vectors, with Polq playing a crucial role^24,25^, we only observed a reduction in AAV integration following the loss of Lig4 (Fig. 7k). However, Lig4 loss also led to increased frequency of Endo-LI, suggesting that Lig4 has distinct roles in exogenous DNA integration and Endo-LI formation and these processes might occur through different mechanisms. Based on these findings, our work suggests that enhancing NHEJ activity—particularly in the absence of AAV as a vector—represents a viable strategy to minimize the formation of both LDs and Endo-LIs.

### Potential strategies to improve genome editing precision *in vivo*

Given that on-target byproducts arise from the repair of DSBs, CRISPR-based technologies that do not induce DSBs, such as base editing and prime editing^1,19^, are likely to avoid such unintended genomic alterations. However, these tools are typically limited to correct small-scale or single-point mutations and often require larger editing reagents, posing significant delivery challenges^22^. Consequently, CRISPR-Cas9, despite its DSB dependency, is expected to continue expanding in clinical applications. Minimizing DSB-induced genomic aberrations is therefore crucial for reducing potential genotoxicity in patients. Our findings provide actionable insights for improving the safety of CRISPR/Cas9-mediated *in vivo* therapies. First, whenever feasible, non-viral delivery methods such as LNP-mediated Cas9 mRNA delivery should be prioritized to avoid the integration of exogenous DNA vectors and reduce off-target effects^28^. Second, elaborate design of sgRNAs by selecting target sites with specific sequence context could increase the likelihood of generating staggered DSBs, which are associated with more predictable repair outcomes^3^. Third, co-delivering small-molecule drugs that enhance NHEJ activity could help to reduce the occurrence of LDs and Endo-LIs, confining edits to more precise DNA regions. Combining these strategies with the use of high-fidelity Cas9 variants to reduce off-target effects could significantly improve the safety of CRISPR/Cas9-based therapies *in vivo*.

We should point out the limitations of our study. First, short-read NGS sequencing was employed to simultaneously investigate the patterns of CRISPR/Cas9-induced indels and large on-target byproducts. While it is currently the gold standard method for quantifying small indels with a low error rate (approximately 0.1%)^10,13^, its sequencing length is limited to 300 bp, making it unsuitable for accurately quantifying large genomic events. As a result, the prevalence of CRISPR/Cas9-induced large on-target byproducts, especially LDs, may be significantly underestimated. Long-read sequencing tools could be considered to yield more accurate detection and characterization of these large byproducts^13,23^. Secondly, although such large on-target modifications were detected, their functional consequence remains unexplored; it is unknown whether they could compromise cellular functionality or promote tumorigenesis. As CRISPR/Cas9-based therapies are already advancing into clinical trials, there is an urgent need for comprehensive safety assessments to balance the risks of gene editing with the pressing demand for treating serious diseases. Lastly, our study was conducted exclusively in MuSCs, it remains unclear whether the findings are applicable to other non-dividing or terminally differentiated cells *in vivo*. Further investigation across diverse cellular states will be necessary to have a comprehensive understanding of the CRISPR/Cas9-mediated *in vivo* editing outcomes.

## Methods

### Mice

All animal experiments were conducted in accordance with the guidelines for the use of laboratory animals established by the Chinese University of Hong Kong (CUHK) and were approved by the Animal Experimentation Ethics Committee (AEEC) of CUHK. Mice were housed in the CUHK Animal Facility under standard conditions with a 12-hour light/12-hour dark cycle. The Cre dependent *Rosa26*^Cas9-EGFP^ knockin mice (B6;129-Gt(ROSA)26Sor^tm1(CAG-cas9*-EGFP^) Fezh/J; stock number 024857) were obtained from the Jackson Laboratory. To generate Cas9 knockin mice, homozygous Pax7^Cre^ mice were crossed with Rosa26^Cas9-EGFP^ mice, as previously described^31,32,34^.

### AAV9 virus production, purification, and injection

AAV9 virus was produced as previously described^31,32^. Briefly, HEK293FT cells were seeded in T75 flasks and transfected at 80-90% confluency with a 1:1:2 ratio of AAV9-sgRNA vector (5 μg), AAV9 serotype plasmid (5 μg), and pDF6 helper plasmid (10 μg) using polyethyleneimine. After 24 hours, the cells were switched to growth medium (DMEM supplemented with FBS, 100 U/mL penicillin, 100 μg/mL streptomycin, and 2 mM L-glutamine) for an additional 48 hours of culture. Cells were then harvested and washed twice with sterile phosphate-buffered saline (PBS). To release the AAV9 virus, the cell pellet was resuspended in AAV lysis buffer [50 mM Tris-HCl (pH 8.0) and 150 mM NaCl] and subjected to three sequential freeze-thaw cycles (liquid nitrogen/37°C). The lysates were treated with Benzonase (Sigma-Aldrich) along with MgCl_2_ (final concentration of 1.6 mM) at 37°C for 30 minutes, followed by centrifugation at 3,000 rpm for 10 minutes. The supernatant was mixed with 1/4 volume of a 5× PEG/NaCl solution [40% PEG8000 (w/v); 2.5 M NaCl] to precipitate the virus at 4°C overnight. After centrifugation at 4,000 g for 30 minutes at 4°C, the pellet was resuspended in sterile PBS and centrifuged again at 3,000g for 10 minutes. The supernatant was filtered through a 0.22 μm sterile filter and passed through a 100 kDa molecular weight cutoff filter (Millipore). The concentrated solution was washed three times with sterile PBS. The titer of the AAV9 virus was determined by quantitative reverse transcription polymerase chain reaction (qRT-PCR) using primers targeting the CMV promoter. The sequences of the primers used are listed in Supplementary Table 9. To profile CRISPR/Cas9 induced editing patterns, 5× 10^11^vg of AAV9-sgRNA or AAV9-dual sgRNA was intramuscularly injected into heterozygous Pax7^Cas9^ mice at P10. To assess the effect of AAV dosage on AAV integration, 1×10^12^ vg, 5×10^11^ vg or 1×10^11^ vg of AAV9-dual sgRNA virus targeting the D16 site were injected intramuscularly into Pax7^Cas9^ mice at P10. To evaluate the effects of NHEJ/MMEJ on the formation of CRISPR/Cas9 induced large on-target modifications, 8×10^11^ vg of AAV9-dual sgRNA virus targeting the CDS regions of Ku80, Lig4, Nbn, and Polq was injected intramuscularly into Pax7^Cas9^ mice at P5, respectively. For the control group, the same dose of AAV9 virus containing the pAAV9-sgRNA backbone without any sgRNA insertion was injected. Then, 5×10^11^ vg of AAV9-single sgRNA was administered at P10 to target the S29 site. MuSCs were isolated four weeks after the AAV injection.

### MuSC isolation by fluorescence-activated cell sorting

MuSCs were isolated by fluorescence-activated cell sorting (FACS) based on established protocols^45^. Briefly, hindlimb muscles from Pax7^Cas9^ mice were collected and digested with collagenase II (1000 U/ml; Worthington) for 90 minutes at 37°C. The tissue suspension was then washed in washing medium [Ham’s F-10 medium (Sigma-Aldrich), 10% heat-inactivated horse serum (HIHS) (Gibco), penicillin/streptomycin (1×, Gibco)]. MuSCs were further liberated by collagenase II (1000 U/ml) and dispase (11 U/ml) treatment for 40 minutes at 37°C. Mononuclear cells were filtered through a 40 μm cell strainer, and GFP+ MuSCs were sorted using a BD FACSAria Fusion cell sorter (BD Biosciences).

### Cell culture

Human embryonic kidney (HEK) 293FT cells (ATCC, CRL-3216) were cultured in Dulbecco’s modified Eagle medium (DMEM) supplemented with 10% fetal bovine serum (FBS), 100 U/mL penicillin, and 100 μg/mL streptomycin in a 5% CO_2_ humidified incubator at 37°C.

### Plasmids

Site-specific sgRNAs were designed using the CRISPOR web tool (http://crispor.tefor.net/)^46^. Oligonucleotides encoding the guide sequences were synthesized and cloned into the AAV9-sgRNA transfer vector [AAV: ITR-U6-sgRNA(backbone)-CMV-DsRed-WPRE-hGHpA-ITR] via the Sap I site. For dual-sgRNA plasmid generation, the second sgRNA cassette containing the U6 promoter and guide sequence was inserted into the AAV9-sgRNA vector using Xba I and Kpn I restriction sites. All sgRNA sequences are provided in Supplementary Table 9.

### Western blotting

Western blotting was performed as previously described^31,34,47,48^. The following dilutions of antibodies were used for each antibody: Histone H3 (Santa Cruz Biotechnology, sc-517576, 1:3000), α-Tubulin (Santa Cruz Biotechnology, sc-23948, 1:2000), Ku80 (Proteintech, 16389-1-AP, 1:1000), Lig4 (Cell Signaling Technology, 14649S, 1:1000), Nbn (Cell Signaling Technology, 3002S, 1:1000), Polq (Cell Signaling Technology, 48160S, 1:1000).

### Targeted deep sequencing

Targeted deep sequencing was performed as previously described^31^. Briefly, genomic DNA from MuSCs infected with AAV9-sgRNA or AAV9-dual sgRNA was amplified using the Q5 High-Fidelity 2× Master Mix (NEB). PCR products were purified using the QIAquick PCR Purification Kit (Qiagen) and subjected to library preparation with the NEBNext® Ultra™ II DNA Library Preparation Kit (NEB). Barcoded libraries were pooled and sequenced on the Illumina HiSeq 1500 platform. The primers used for Deep-seq are listed in Supplementary Table 1.

### Variant detection for single-sgRNA target sites

All sequencing data for single-sgRNA groups were generated in-house, except for samples S12, which were sourced from published paper^31^. In house scripts were developed to extract NGS reads containing target-specific primer sequences. Quality filtering was performed using Trimmomatic (v0.39)^49^ with the parameter AVGQUAL:30 to eliminate low-quality reads. In instances where paired reads exhibited significant quality disparities, only the higher-quality end was retained. The processed reads were then analyzed using SIQ (v1.3)^36^ through a two-stage workflow: (i) merging overlapping paired-end reads and (ii) aligning the merged consensus sequences to target references for variant detection, utilizing default parameters (Min support: 2; Max base error: 0.08; TINS search distance: 100). While most analyses relied on paired-end data, single-end processing was employed in three specific scenarios: (i) when targets exceeded 300 bp, (ii) in cases of insufficient merge efficiency, or (iii) due to substantial quality discordance between read pairs (Supplementary Table 2). The outcomes were classified into the following categories: wild-type (WT), single-nucleotide variant (SNV), insertion, deletion, delins (an insertion coinciding with a deletion), TINS (a deletion accompanied by an insertion sourced from the adjacent sequence within 100 bp of the cut site), tandem duplication (TD), and TD compound (a TD that incorporates additional insertions). For each event, the fraction, position, and size of indels, as well as the MH for deletions were provided. In terms of relative mutation positioning, SIQ defines the canonical cut site as position 0, with positions to the left of the cut site assigned negative values and those to the right assigned positive values. Given that targets vary in length, we ensured a fair evaluation of on-target editing by defining a 60 bp window centered on the canonical cut site for all target sites. Events overlapping with this window (delRelativeStart ≤ 30 and delRelativeEnd ≥ -30) were included in the subsequent analysis, with their corresponding fractions adjusted accordingly. Indel events—including insertion, deletion, delins, TINS, TD, and TD compound—were classified as edited events, while SNVs were categorized as WT events. Finally, mutational spectra were visualized using tornado plots generated via the SIQPlotteR website (https://siq.researchlumc.nl/SIQPlotteR/). Target genomic locations were displayed using the R package “RIdeogram”^50^.

### Definition and quantitative analysis of editing precision

The precision at a target site was defined by the average frequency of the most common indel among total edited events across two biological replicates. All targets were subsequently categorized into three groups based on precision: imprecise (≤ 0.25), medium (0.25 < middle ≤ 0.5), and precise (> 0.5)^38^. To assess the number of indel types within each precision group, only events with frequencies in total edited events exceeding 0.005 were included. This approach was taken to reduce bias resulting from differences in sequencing depths. The number of indel types for each target was averaged across two biological replicates. To evaluate indel size within each precision group, deletions were defined as negative values and insertions as positive. The absolute indel size was calculated as |delSize + insSize|. These sizes were weighted by their frequencies in total edited events and summed up to obtain the final indel size for each replicate. Subsequently, the final indel sizes were averaged across two biological replicates for each target. Statistical significance was assessed using a t-test.

### Processing and correlation analysis of epigenomic features

ChIP-Seq data was obtained from the Genome Sequence Archive (GSA) under accession CRA002490^42^. ATAC-Seq data was downloaded from the Gene Expression Omnibus (GEO) under accession number GSM1552830^51^. Detailed information is provided in Supplementary Table 8. ChIP-Seq reads were trimmed using Trimmomatic (v0.36)^49^ and aligned to the mm10 genome using Bowtie 2 (v2.4.4)^52^. Duplicate reads were removed using Picard (v2.26.0)^53^. Peak calling was performed using MACS2 (v2.2.6)^8^, with a control file (Supplementary Table 8). Finally, IDR (v2.0.0)^54^ was used to combine replicates and filter for highly reproducible peaks. ATAC-Seq reads were trimmed using Trim Galore (v0.6.10) (https://github.com/FelixKrueger/TrimGalore) and aligned to the mm10 genome using BWA (v0.7.17-r1188)^55^. Reads with a mapping quality <30 and duplicate reads were removed using SAMtools (v1.5)^56^ and Sambamba (v1.0.0)^57^. To improve peak-calling accuracy, blacklisted regions (ENCODE mm10 v2)^58^ were removed using BEDTools^59^, followed by Tn5 offsets correction with alignmentSieve (deepTools v3.5.5)^60^. Peaks were called without a control file using the ChIP-Seq pipeline. To calculate the average signal intensity over target regions, only replicate 1 of epigenomic data was used (Supplementary Table 8). The BAM files were converted to BigWig format using BamCoverage (v3.5.5) ^60^. Next, bigWigAverageOverBed (v2)^61^ was employed to calculate the average intensity over bases, treating non-covered bases as zeroes in each target. Spearman’s correlation was calculated between the average intensity of each target and its corresponding average TIF across replicates for all targets. Correlations of average intensity with other variables were performed similarly.

### Quantitative and directional analysis of MH-mediated deletions

The expected ratio for each k-mer size in MH was calculated as 0.25^k^. The average deletion size for k-bp MH deletions was determined by weighting the sizes of the k-bp MH deletions based on their frequencies among the total k-bp MH deletions and summing these values to obtain a final size. The final k-bp MH deletion sizes were then averaged across two biological replicates for each target. Statistical significance was evaluated using a t-test.

SIQ designated position 0 as the canonical cut site, with negative values assigned to left-side positions and positive values to right-side positions. In directional analysis, deletions were classified into downstream (within the PAM-containing flank), upstream (opposite flank), or spanning (crossing the cut site).

### Resolution of TIs at single-sgRNA target sites

The frequency of TIs in k-bp insertions without deletions at the cut site (SIQ-reported relative start position 0) was determined by aligning the inserted sequences to reference segments of equal length. These segments extended either (i) rightward from position -3 or (ii) leftward from position -4 relative to the NGG PAM (Fig. 3a). When inserted nucleotides generate or occur within repetitive sequences at the cut site (e.g., a “C” adjacent to another “C” forming “CC,” or a “G” extending “GG” to “GGG”), the exact insertion position becomes ambiguous. Consequently, SIQ may assign relative start positions as 0, 1, or 2. To address this uncertainty, these events were manually curated. For the allele frequency heatmap visualization, both reference and inserted alleles were standardized to the NGG PAM orientation.

### Variant detection for dual-sgRNA target sites and characterization of editing outcomes

All sequencing data for dual-sgRNA groups were generated in-house, except for samples D5 and D32, which were sourced from published paper^31^. For data analysis, primer-specific reads were extracted, and quality control was performed on the NGS data for each dual-sgRNA site (Supplementary Table 1), following the same workflow applied to single-sgRNA sites. Next, a two-stage alignment approach, adapted from Chakrabarti et al^38^, was implemented. In the initial stage, reads were aligned to the reference genome using BBMap (v39.01)^62^. During this phase, reads that aligned as proper pairs were discarded, and only the remaining reads were retained for further analysis using Samtools (v1.6)^56^. In the second stage, these remaining reads were aligned to the reference genome using BBMap (v39.01)^62^ with a threshold that permitted indels of up to 3000 bp. Finally, unmapped reads and secondary alignments were filtered out utilizing Samtools (v1.6)^56^ and Sambamba (v1.0.0)^57^. Indels were detected and characterized using R scripts adapted from Chakrabarti et al.,^38^ with positions and lengths determined through CIGAR string parsing. For each target’s two cut sites (3 bp upstream of NGG PAMs), indels overlapping the ±10 bp window surrounding either cut site were analyzed. Reads containing these indels were classified as edited reads and were categorized into three types: type 1, which includes indels spanning the windows of both cut sites; type 2, consisting of indels that overlap the window of only one cut site; and type 3, involving concurrent indels that overlap the windows of both cut sites without spanning. Type 1 is prioritized over the other types. Unedited reads were defined as: (i) quality-trimmed reads before alignment, excluding the remaining reads from the initial alignment, and (ii) reads whose indels fell completely outside both cut site windows. For type 1 analysis, type 1 reads were extracted to SIQ (v1.3)^36^ and precisely ligated sequences were utilized as references to explore the accurate ligation (Supplementary Table 2). For subsequent analysis, only targets with a total event count exceeding 100 were considered. SNVs and WT were classified as accurate events, and PAM orientations were assigned as ’W’ for NGG and ’C’ for CCN. For TI analysis, 1 bp TIs were defined as insertions matching either the -3 or -4 nucleotide upstream of either PAM. 2 bp TIs were defined as insertions that matched either the -3:-2 or -5:-4 dinucleotides of either PAM, or any of the four combinations consisting of one nucleotide from the -3 or -4 position of PAM1 and one nucleotide from the -3 or -4 position of PAM2.

### Comprehensive analysis of the origins and characteristics of LIs

Parallel alignments were conducted using BWA (v0.7.17-r1188)^55^ to the AAV vector sequence and the mm10 reference genome. LIs were processed through distinct alignment pipelines based on size: (i) for LIs ≤ 70 bp, bwa-aln was used followed by bwa-samse, and (ii) for LIs > 70 bp, bwa-mem was employed. All alignments were filtered using Samtools (v1.6)^56^ to eliminate unmapped reads and secondary alignments. When LIs were simultaneously mapped to both the mm10 and AAV sequences, the origin from mm10 took priority over that from AAV.

For functional annotations of Endo-LIs, genomic locations of Endo-LI donors were annotated using the R package “ChIPseeker”^63,64^, with those within ± 3 kb of transcription start sites categorized as promoters. En-LI donor locations overlapping with H3K27ac peaks were designated as enhancers (Supplementary Table 8), with enhancers prioritized over other functional categories. The downstream category was classified as intergenic. For repetitive element annotation, the RepeatMasker output for the mm10 assembly was combined with the mm10 gap file, both obtained from the UCSC Genome Browser. Centromeres and satellites were classified together as centromeres. LINE, LTR, and SINE elements were grouped as retrotransposons. To resolve overlaps, the following priority hierarchy was employed: retrotransposons > transposons > centromeres > simple repeats > other repeats.

To analyze MH usage in Endo-LIs/AAV-LIs, custom R scripts were developed for detection. For each LI, the acceptor and donor site locations were identified within their respective reference genomes. Extended sequences flanking both sides of these sites were then extracted. Subsequently, the sequence consistency between the donor and acceptor were assessed by comparing extensions ranging from 1 to 15 nucleotides.

For analyzing LIs within TAD, the processed .hic file of WT proliferating myoblasts from GEO accession GSE157339^42^ was used. TADs were identified at a 40k resolution using Juicer Tools (v2.20.00.ac)^65^ with the arrowhead method and the -k KR parameter. Since the .hic file was based on mm9, LiftOver^66^ was utilized to convert the TAD locations to mm10. In our analysis, only LIs with donor and acceptor located on the same chromosome were considered. The intersect command from BEDTools (v2.27.1)^59^ was used to determine whether the donor and acceptor were situated within consistent TADs, followed by a paired t-test to evaluate significance. The visualization of TAD heatmap was generated using the mm9 assembly.

### Comparison of LD, LI, and Endo-LI type counts across precision groups and JSS analysis

To reduce sequencing depth bias, only LD, LI, and Endo-LI types constituting at least 0.01% of total indels and originating from targets with more than 10,000 total indels in both replicates were included. The counts of these LD, LI, and Endo-LI types were then compared across the I, M, and P groups in Fig. 5f, 5s and 5t. For JSS analyses in Fig. 5b, 5i and 5l, all LD, LI, and Endo-LI types meeting these criteria were compared between samples.

### Evaluation of prediction tools on *in vivo* repair outcomes

Three commonly used prediction tools were employed: inDelPhi^16^, ForeCAST^17^, and Lindel^8^. For inDelPhi, parameters were set to mESC cell type, 0.05% minimal frequency, and default indel length range. The complete target sequences used in SIQ were used as the input data for the analyses of inDelPhi and ForeCAST. For Lindel, the input consisted of 65 bp, including 30 bp upstream and 35 bp downstream of the cleavage site. Indels were classified into five categories: 1 bp insertions, > 1 bp insertions, 1 bp deletions, 2 bp deletions, and ≥ 3 bp deletions. Predictions were statistically compared to SIQ measurements (averaged across two replicates per target) using t-tests. Pearson correlation analysis performed on vectorized representations of the five indel categories for each target.

### Data availability

All Deep-seq data generated in this study have been deposited in Gene Expression Omnibus (GEO) database under the accession code GSE303540.

## Author contributions

Conceived and designed the experiments: H.W., H.S., L.H. and Y.F. Performed the experiments: L.H., Z.W. and Q.Z. Analyzed the data: Y.F. Wrote the paper: H.W., L.H. and Y.F. Reviewed and edited the manuscript: H.W., H.S., L.H. and Y.F.

## Funding

This work was supported by National Key R&D Program of China to H.W. (project code: 2022YFA0806003); Non-Communicable Chronic Disease-National Science and Technology Major Project of China to H. W. (project code: 2024ZD0530400); The InnoHK initiative of the Innovation and Technology Commission of the Hong Kong Special Administrative Region Government to H.W. (project code:INNOHK22SC01); Health and Medical Research Fund (HMRF) from Health Bureau of HK to H.W. (project codes: 10210906 and 08190626); Theme-based Research Scheme (TRS) from RGC to H.W. (project code:T13-602/21-N);General Research Fund (GRF) from Research Grants Council (RGC) of the Hong Kong Special Administrative Region, China to H.W. (project codes: 14108225, 14106521, 14105123, 14103522, and 14105823 to H.W.); the National Natural Science Foundation of China (NSFC) to H.W. (project codes: 82172436); Area of Excellence Scheme (AoE) from RGC to H.W. (project code: AoE/M-402/20). The Chinese University of Hong Kong (CUHK) Strategic Seed Funding for Collaborative Research Scheme (SSFCRS) to H.W.

### Conflict of interest statement

The authors declare that they have no conflict of interest.

## Notes

### Competing Interest Statement

The authors have declared no competing interest.

