## Supplementary information for "*In vivo* systematic detection of the outcomes of CRISPR/Cas9 mediated DNA repair in skeletal muscle stem cells"

**Supplementary Materials**

**List of Extended Data Figures**

Extended Data Figure 1. *In vivo* mapping of CRISPR/Cas9 induced indel profiles in MuSCs using single-sgRNAs.

Extended Data Figure 2. Impact of chromatin states on editing precision.

Extended Data Figure 3. Extensive use of microhomology in CRISPR/Cas9 induced deletions in MuSCs *in vivo*.

Extended Data Figure 4. Limited predicting power of the machine learning models *in vivo*.

Extended Data Figure 5. *In vivo* mapping of CRISPR/Cas9 induced indel profiles in MuSCs using dual-sgRNAs.

Extended Data Figure 6. Extensive use of microhomology in dual sgRNAs induced deletion in MuSCs *in vivo*.

Extended Data Figure 7. Analysis of CRISPR/Cas9 induced large on-target genomic modifications by single-sgRNAs in MuSCs *in vivo*

Extended Data Figure 8. Analysis of CRISPR/Cas9 induced large on-target genomic modifications by dual-sgRNAs in MuSCs *in vivo*

Extended Data Figure 9. AAV integration is a general outcome of CRISPR/Cas9-induced editing in MuSCs *in vivo*.

Extended Data Figure 10. *In vivo* perturbation of NHEJ/MMEJ pathways can modulate CRISPR/Cas9 editing in MuSCs

**List of Supplementary Tables**

Supplementary Table 1. List of single and dual sgRNA target sites.

Supplementary Table 2. SIQ run information.

Supplementary Table 3. SIQ outcomes for single-sgRNA sites.

Supplementary Table 4. Jaccard similarity scores between target sites.

Supplementary Table 5. SIQ outcomes for dual-sgRNA sites.

Supplementary Table 6. SIQ outcomes for AAV dose groups.

Supplementary Table 7. SIQ outcomes for *in vivo* perturbation.

Supplementary Table 8. Sources of ChIP-seq and ATAC-seq data used in the study.

Supplementary Table 9. Sequences of oligos used in this study.

### He L and Fu Y et. al. Extended Data Figure 1

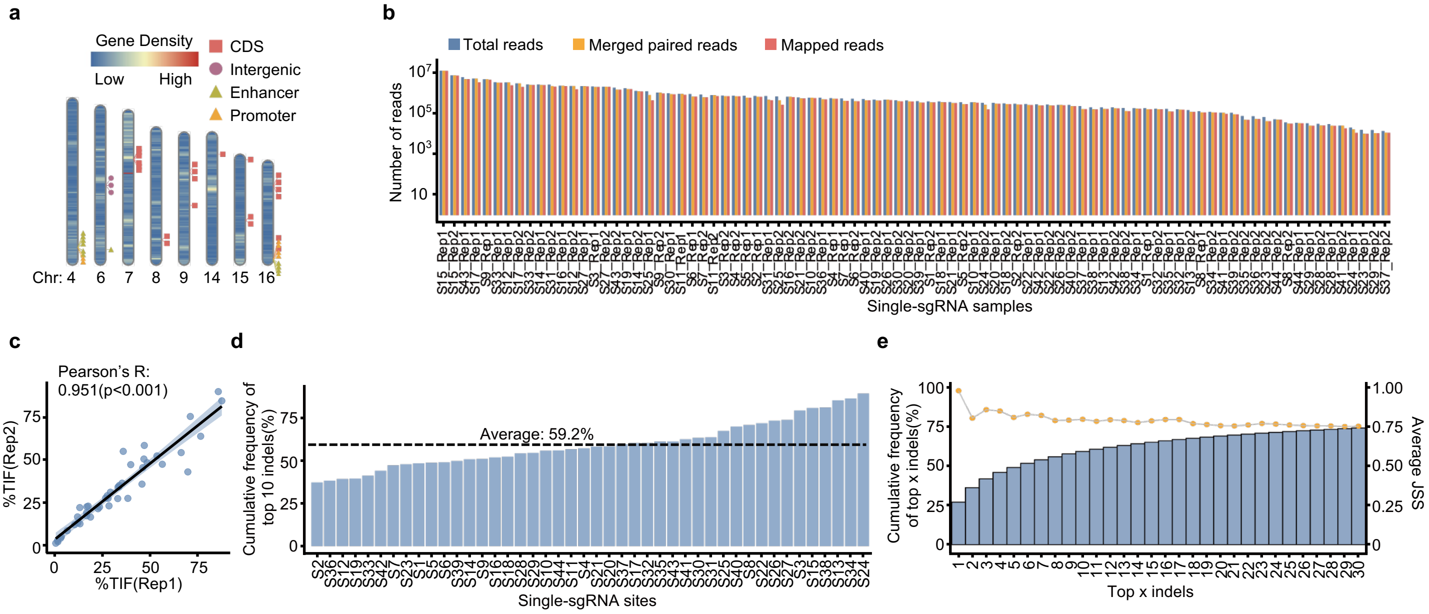

**Extended Data Figure 1. *In vivo* mapping of CRISPR/Cas9 induced indel profiles in MuSCs using single-sgRNAs.** (**a**) Distribution of the 44 single-sgRNA targets across chromosomes. Genomic annotation of each target is shown. (**b**) Sequences encompassing each target site were amplified and subjected to deep-sequencing. Sequencing reads alignment metrics for the 44 targets (each with 2 biological replicates) are shown. (**c**) Correlation of TIF between two biological replicates across all 44 targets. (**d**) The cumulative indel frequency of the top ten most frequent indels for each target. The average frequency across all sites is shown by the dash line. (**e**) The cumulative indel frequency of the top 1 to 30 most frequent indels across 44 targets, along with the corresponding average JSS of paired replicates across all targets.

### He L and Fu Y et. al. Extended Data Figure 2

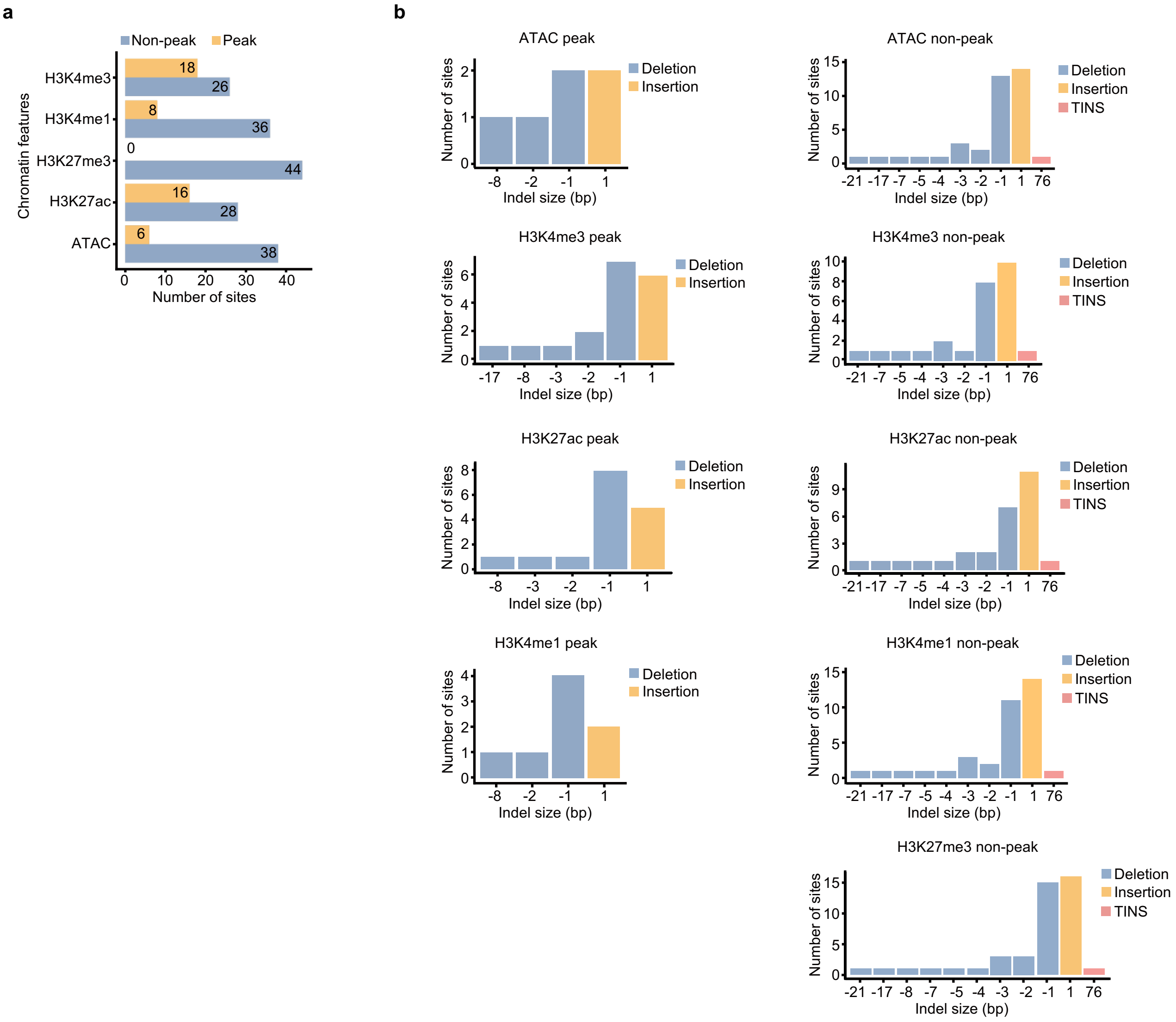

**Extended Data Figure 2. Impact of chromatin states on editing precision.** (**a**) Numbers of sites located in peak or non-peak regions of each indicated chromatin feature. (**b**) The number of sites with the indicated type of top 1 indels in peak or non-peak regions of each indicated chromatin feature.

### He L and Fu Y et. al. Extended Data Figure 3

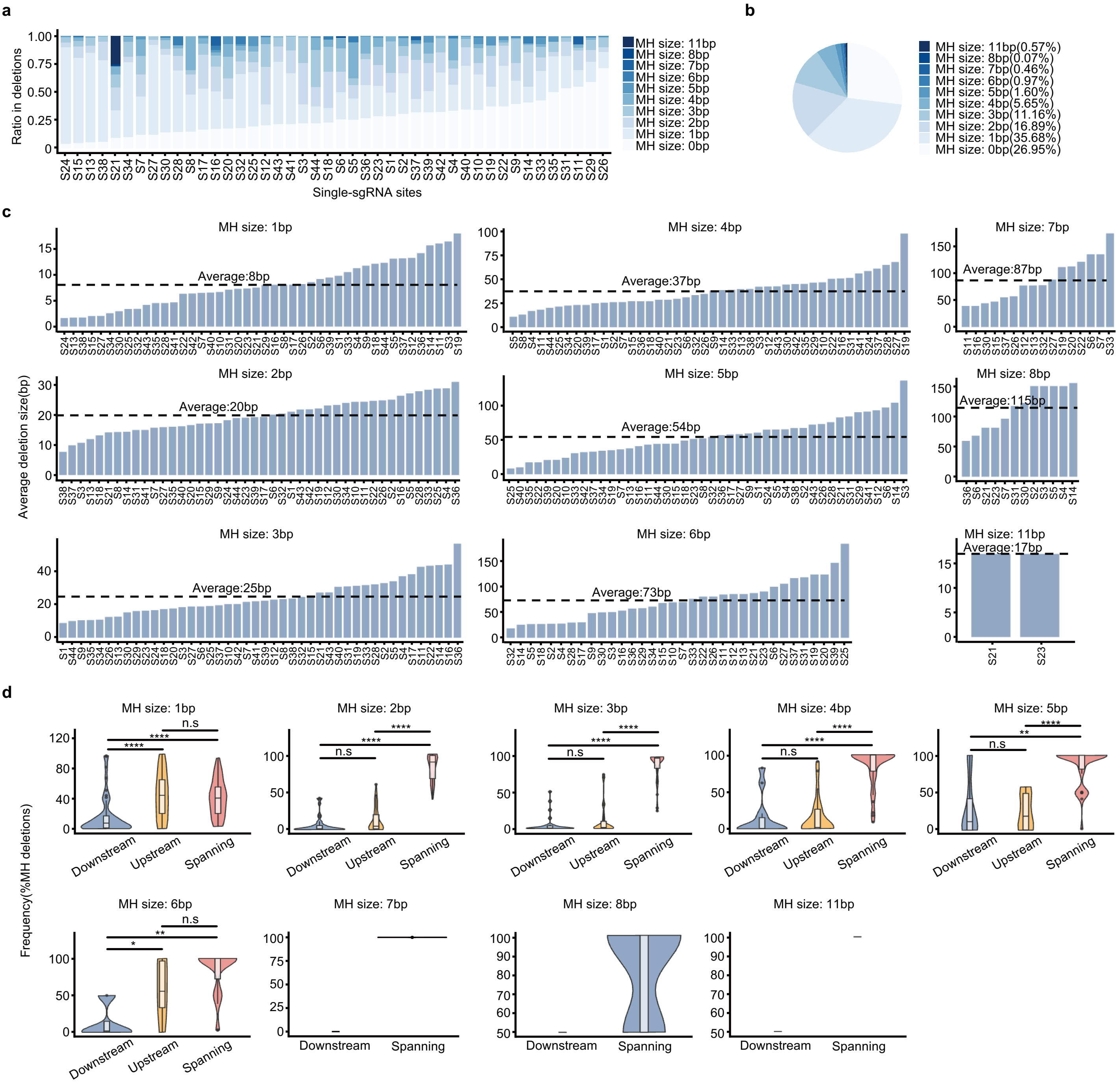

**Extended Data Figure 3. Extensive use of microhomology in CRISPR/Cas9 induced deletions in MuSCs *in vivo*.** (**a**) Ratio of deletions with MH of varying sizes at each site. (**b**) Pie chart showing the average frequency of deletions with MH of varying sizes across all sites. (**c**) The average deletion sizes categorized by varying MH lengths. (**d**) Violin plots illustrating the frequency of deletions with varying MH sizes categorized by direction. Statistical significance was assessed using a t-test. *$p < 0.05$, ** $p < 0.01$, ****$p < 0.0001$ and ns, no significance.

### He L and Fu Y et. al. Extended Data Figure 4

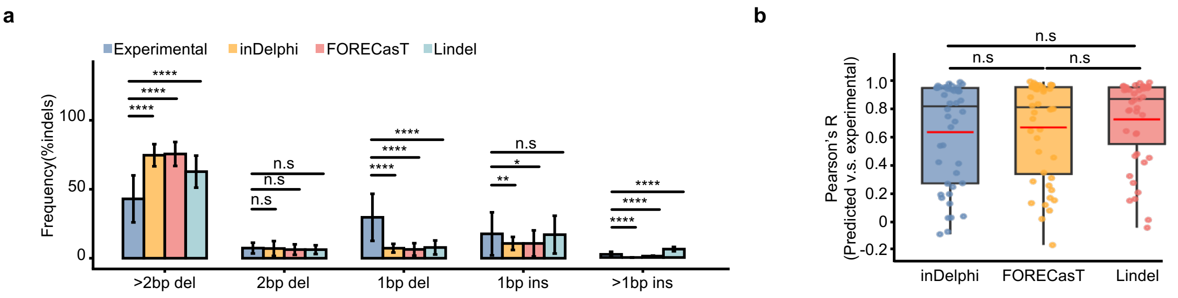

**Extended Data Figure 4. Limited predicting power of the machine learning models *in vivo*.** (**a**) Comparison between the frequency in total indels predicted by inDelphi, FORECasT, and Lindel and the experimentally measured for > 2 bp deletions, 2 bp deletions, 1 bp deletions, 1 bp insertions and > 1 bp insertions across the 44 sites, respectively. (**b**) Pearson coefficients were calculated for the predicted vs. experimentally measured indel frequency for each target. Statistical significance for (**a**) and (**b**) was calculated using a t-test. *$p < 0.05$, ** $p < 0.01$, ****$p < 0.0001$ and ns, no significance.

### He L and Fu Y et. al. Extended Data Figure 5

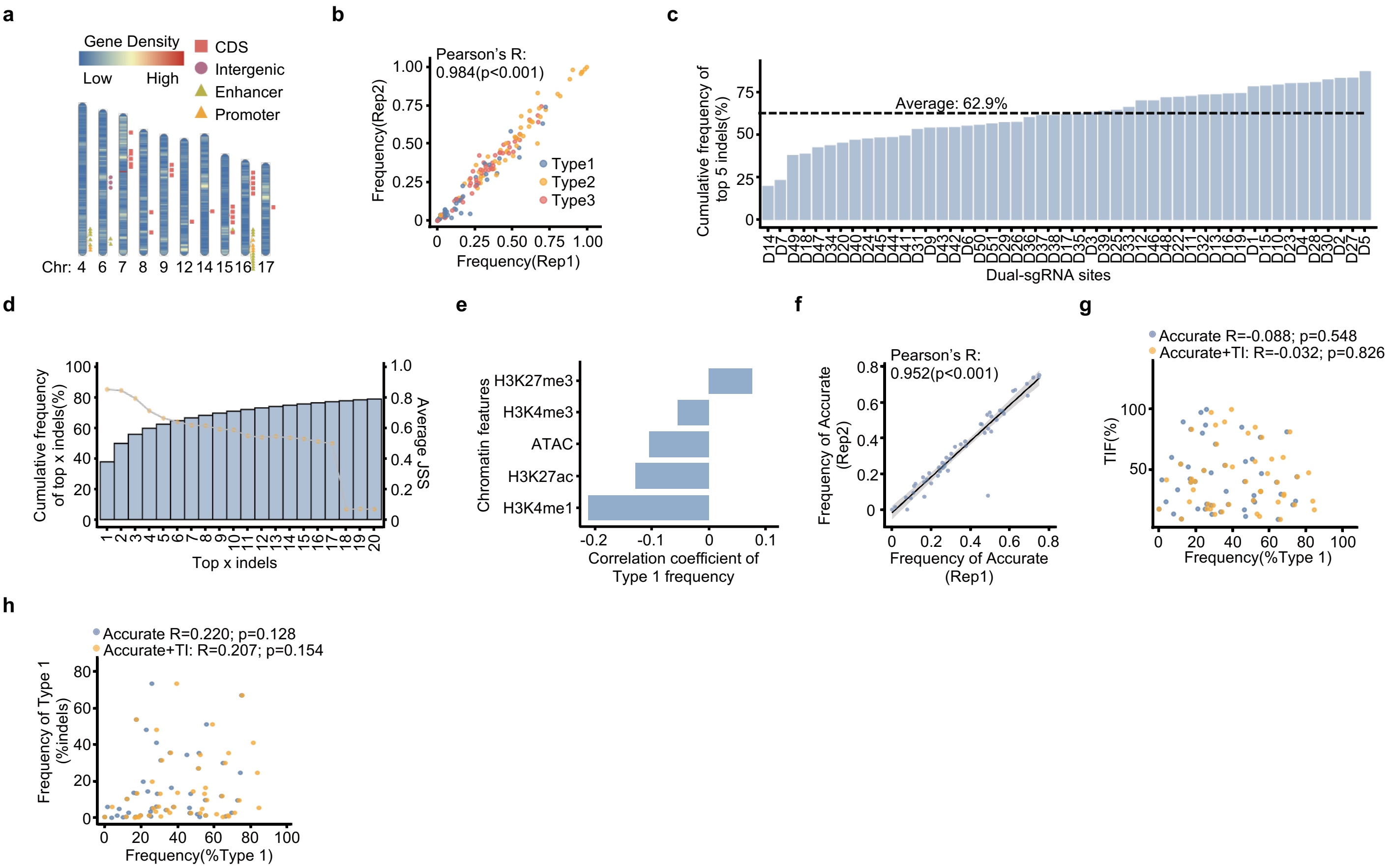

**Extended Data Figure 5. *In vivo* mapping of CRISPR/Cas9 induced indel profiles in MuSCs using dual-sgRNAs.** (**a**) Distribution of the 51 dual-sgRNA targets across chromosomes. Genomic annotation of each target is shown. (**b**) Correlation of Type 1, Type 2 and Type 3 indel frequency between two biological replicates across all 51 targets. (**c**) The cumulative indel frequency of the top five most frequent indels for each site. The average cumulative indel frequency across all sites is shown. (**d**) The cumulative indel frequency of the top 1 to 20 most frequent indels across 51 targets, along with the corresponding average JSS of paired replicates across all targets. (**e**) Spearman's correlation between Type 1 frequency and indicated chromatin features. (**f**) Pearson's correlation of the frequency of accurate ligation (Accurate) in Type 1 events between two biological replicates across targets. (**g**) Correlation of TIF with the frequency of Accurate (blue) or Accurate + TI (orange) in Type 1 indels. (**h**) Correlation of Type 1 frequency with the frequency of Accurate or Accurate + TI in Type 1 indels.

### He L and Fu Y et. al. Extended Data Figure 6

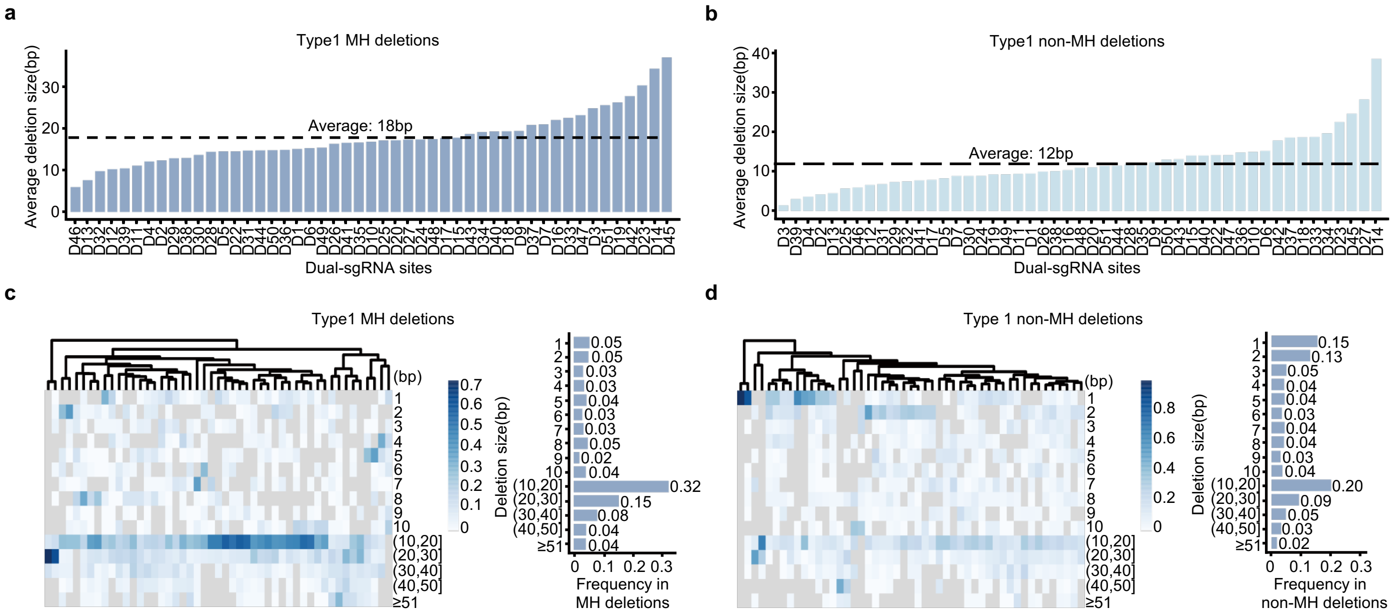

**Extended Data Figure 6. Extensive use of microhomology in dual sgRNAs induced deletion in MuSCs *in vivo*.** (**a**-**b**) Average deletion size for MH (**a**) and non-MH (**b**) mediated deletions for each site. The average deletion size for MH and non-MH mediated deletions across all dual-sgRNA sites is shown by dash line. (**c**) Left: Heatmaps illustrating the frequency of MH mediated deletions categorized by size across dual-sgRNA targets. The targets are clustered using the "complete" hierarchical clustering. Right: The average frequency across all targets for each category with varying deletion size in MH mediated deletions. (**d**) The above analysis was conducted in non-MH mediated deletions.

### He L and Fu Y et. al. Extended Data Figure 7

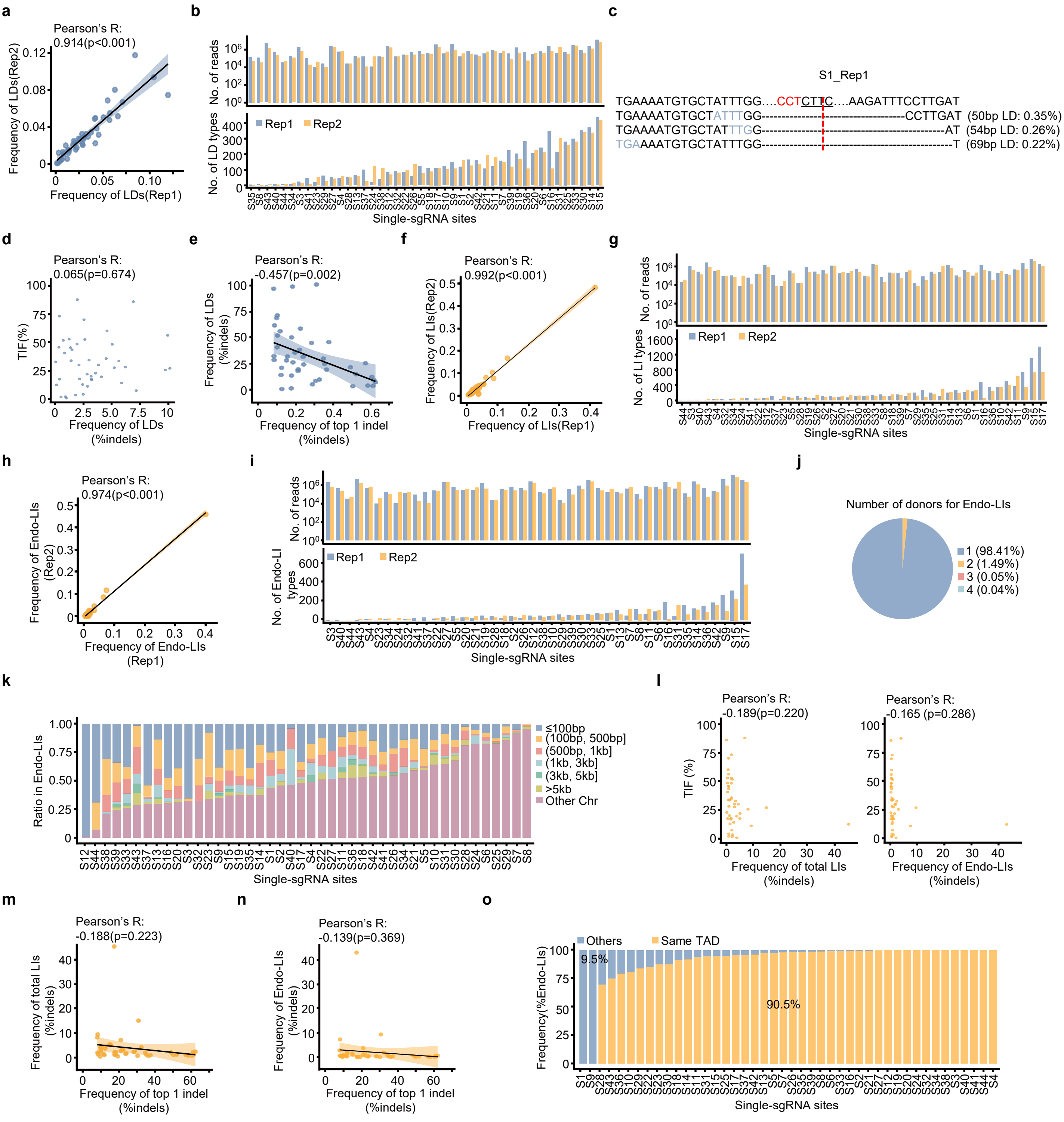

**Extended Data Figure 7. Analysis of CRISPR/Cas9 induced large on-target genomic modifications by single-sgRNAs in MuSCs *in vivo*.** (**a**) Pearson's correlation of LD frequency in total indels between two biological replicates across single-sgRNA targets. (**b**) Number of LD types defined by their start sites and the deleted sequences in each biological replicate across all targets (Lower) and number of sequencing reads for each sample (Upper). (**c**) An example showing the frequency of LDs with MH on S1. The MH tracks are marked as blue. (**d**) Pearson's correlation between TIF and LD frequency. (**e**) Pearson's correlation between LD frequency in all indels and editing precision determined by the frequency of top 1 indel. (**f**) Pearson's correlation of LI frequency in total indels between two biological replicates across single-sgRNA targets. (**g**) Number of LI types defined by their start sites and the inserted sequences in each biological replicate across all targets (Lower) and number of sequencing reads for each sample (Upper). (**h**) Pearson's correlation of Endo-LI frequency between two biological replicates across single-sgRNA targets. (**i**) Number of Endo-LI types defined by their start sites and the inserted sequences in each biological replicate across all targets (Lower) and number of sequencing reads for each sample (Upper). (**j**) Pie charting showing the number distribution of donors for Endo-LIs. (**k**) Distance distribution between insertion donors for Endo-LIs and associated DSBs for each site. (**l**) Pearson's correlation of TIF with total LI (Left) and Endo-LI (Right) frequency. (**m**) Pearson's correlation between total LI frequency and editing precision determined by the frequency of top 1 indel. (**n**) Pearson's correlation between Endo-LI frequency and editing precision determined by the frequency of top 1 indel. (**o**) Frequency of donor sequences originating from the DSB-containing TADs or other loci on the same chromosome for each site.

### He L and Fu Y et. al. Extended Data Figure 8

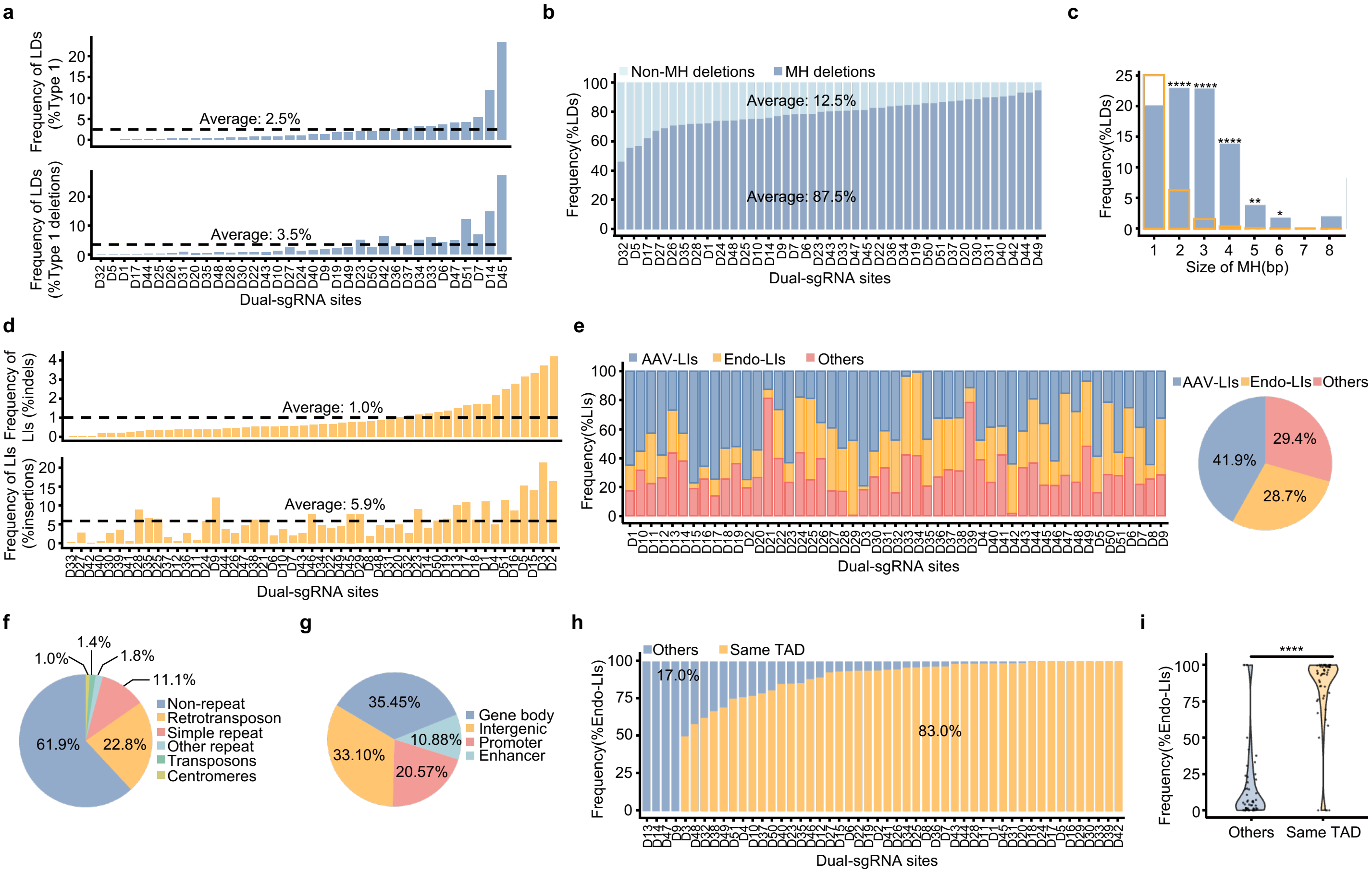

**Extended Data Figure 8. Analysis of CRISPR/Cas9 induced large on-target genomic modifications induced by dual-sgRNAs in MuSCs *in vivo*.** (**a**) Frequency of ≥ 50 bp LDs for each dual-sgRNA site, calculated as percentage in total Type 1 events (Upper) and total Type 1 deletions (Lower). The average frequency of LDs across all sites is shown by dash line. (**b**) Frequency of MH and non-MH mediated LDs for each dual-sgRNA target. The average frequency of MH and non-MH mediated LDs across all sites is shown. (**c**) Frequency of LDs with MH of varying sizes. The orange bar represents the expected frequency for each MH size. (**d**) Frequency of ≥ 20 bp LIs for each dual-sgRNA site, calculated as percentage in total indels (Upper) and total insertions (Lower). The average frequency of LIs is shown by dash line. (**e**) Left: Frequency of different types of LIs for each dual-sgRNA target. Right: Pie chart showing the average frequency of each LI type across all sites. (**f**) Pie chart showing the frequency of donor sequence categorized by types of repetitive elements for Endo-LIs. (**g**) Pie chart showing the frequency of donor sequence categorized by genomic annotations for Endo-LIs. (**h**) Frequency of donor sequences originating from the DSB-containing TADs or other loci on the same chromosome for each dual-sgRNA site. (**i**) Violin plot showing significantly higher frequency of donor sequences originating from DSB-containing TADs compared to other loci on the same chromosome. A one-tail t-test was used for (c) and a paired t-test was used for (**i**). $*p < 0.05$, $** p < 0.01$, $****p < 0.0001$.

### He L and Fu Y et. al. Extended Data Figure 9

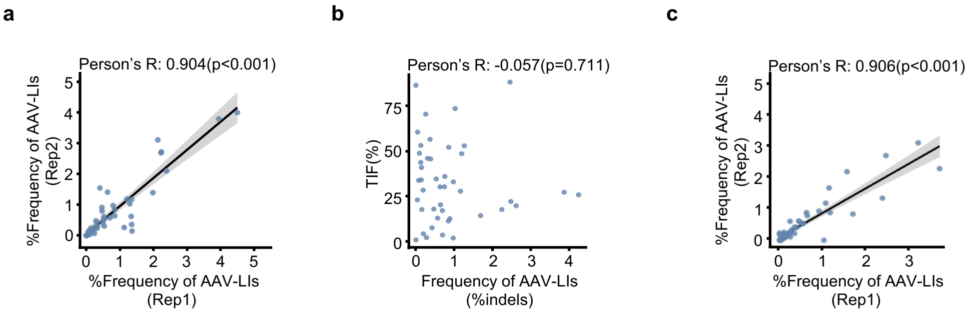

**Extended Data Figure 9. AAV integration is a general outcome of CRISPR/Cas9-induced editing in MuSCs *in vivo*.** (**a**) Pearson's correlation of AAV-LI frequency in total indels between two biological replicates across single-sgRNA targets. (**b**) Pearson's correlation of TIF with AAV-LI frequency. (**c**) Pearson's correlation of AAV-LI frequency between two biological replicates across dual-sgRNA targets.

### He L and Fu Y et. al. Extended Data Figure 10

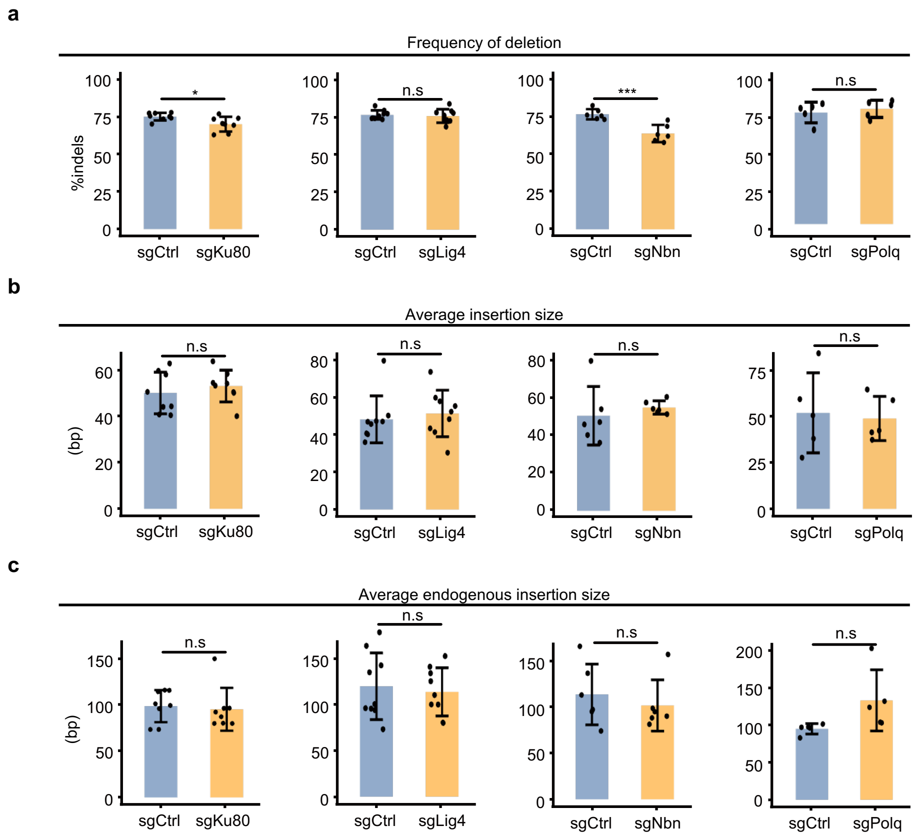

**Extended Data Figure 10. *In vivo* perturbation of NHEJ/MMEJ pathways can modulate CRISPR/Cas9 editing in MuSCs.** (**a**-**c**) Frequency of deletions (**a**), average insertion size (**b**) and average endogenous insertion size (**c**) after *in vivo* depletion of Ku80, Lig4, Nbn or Polq is shown. n=5 to 9 mice in each group. Statistical significance was calculated using a paired t-test for (**a**) - (**c**). $^{*}p < 0.05$, $^{***}p < 0.001$ and ns, no significance.
